# The Neural Code of Neuroticism

**DOI:** 10.64898/2026.01.19.700296

**Authors:** Johanna L. Popp, Martin Weiß, Joshua Faskowitz, Kirsten Hilger

## Abstract

High neuroticism is a risk factor for mental disorders. Understanding whether individuals share a common foundation in brain function underlying neuroticism is therefore an essential goal of neuroscience. We applied Inter-Subject Representational Similarity Analysis on data from 174 Human Connectome Project participants to investigate if their similarity in neuroticism is reflected in their similarity of fMRI-recorded brain activity during movie watching. To test behavioral personality theories that consider trait expression as dependent on situational context, we examined whether brain-trait representational similarity varied between trait-relevant and trait-irrelevant movie scenes (independently rated, *N* = 86). Higher neuroticism was associated with greater heterogeneity in brain responses, a pattern that was particularly pronounced during trait-relevant scenes. Our study informs dimensional conceptualizations of psychopathology viewing neuroticism as risk factor and extends behavioral personality theories to the neural level. Broadly, it highlights the value of naturalistic imaging and underscores the importance of stimulus selection when investigating brain-behavior associations.

## Introduction

Neuroticism is a major dimension of human personality differences^1,2^. It reflects the relatively stable individual tendency to experience negative emotions such as anxiety or depression^3,4^, qualifying it as personality trait^5^. High neuroticism is a risk factor for many mental disorders, including anxiety, mood, and substance use disorders^6–8^. Understanding its neurobiological basis is therefore an essential goal of neuroscience research.

Genetic factors account for 30-60% of the variance in neuroticism^9^, indicating a partial biological foundation. In interaction with environmental influences, this genetic basis contributes to variability in brain structure and function^10^ which is thought to underlie individual differences in this personality trait^3,11^. Neuroimaging studies have identified structural and functional brain correlates of neuroticism primarily in limbic/subcortical (e.g., amygdala, hippocampus) and frontal regions (e.g., dorsolateral prefrontal cortex, anterior cingulate cortex), though findings have been inconsistent^12–14^. Given the broad cognitive and affective implication of neuroticism, whole-brain or network-level analyses might be more appropriate than the restricted focus on predefined regions of interest^15^. Further, functional brain measures linked to neuroticism may be particularly promising, as unlike structural brain measures that reflect temporally stable anatomical characteristics, they can provide a dynamic, context-sensitive perspective on how individuals perceive and respond to the external world. To date, however, most functional MRI (fMRI) research on neuroticism has employed resting-state or standard task paradigms, yielding little consensus on how global brain function relates to this trait^12,14^.

Naturalistic stimuli, such as movies or spoken narratives, offer several advantages over resting-state and traditional task paradigms including a) increased ecological validity^16^, b) improved subject compliance and reduced head motion^17,18^, and c) ease of implementation^19^. Although, in general, naturalistic stimuli evoke highly synchronized spatiotemporal patterns of brain activity across participants^20,21^, significant and reliable individual differences have been shown to persist on top of this shared response^22–24^. Inter-subject correlation (ISC^20,25^) can be used to analyze these differences in brain activity and, by exploiting the time-locked nature of naturalistic stimuli, can ultimately isolate stimulus-driven signal^22^. Although ISCs can, in principle, be related to traits (e.g., via linear mixed-effects model^26^), these approaches do not provide frameworks for validly distinguishing trait-related from common stimulus-driven brain responses, thus limiting critical insight. Inter-subject representational similarity analysis (IS-RSA^22^) enables this distinction, which is realized by comparing between-subject patterns of similarity in brain activity (assessed with ISC) to between-subject patterns of similarity reflected in behavioral attributes, such as personality traits. As inter-subject trait similarity can be determined with different computational models^22^, IS-RSA not only enables detecting whether and where there exists a shared neural foundation of a trait, i.e., inter-subject similarity at the trait level is represented at the brain level (brain-trait representational similarity), but also allows for testing specific assumptions about the nature of the relationship between inter-subject similarity in brain activity, also referred to as inter-subject neural synchrony, and inter-subject similarity in traits.

Prior studies have applied various methods to link inter-subject similarity in brain activity (i.e., ISC) during naturalistic paradigms to trait-like factors including trait-paranoia^26^, working memory^22^, cognitive styles^27^, psychological well-being^28^, socio-sexual desire and self-control preferences^29^, depressive symptoms^30^, and even to personality traits^22,31^. This suggests that naturalistic stimuli can act as ‘implicit primes’, activating trait-relevant neural features, which leads to differential, trait-related, neural responses to identical stimuli^22^. Even though pioneering work has started applying IS-RSA to investigate the neural underpinnings of personality^22,31^, no study specifically focused on neuroticism nor thoroughly evaluated the form of relationship between inter-subject trait similarity and inter-subject similarity in brain activity.

Contemporary personality theories like the Cybernetic Big Five Theory (CB5T)^32^ or the Trait Activation Theory (TAT)^33^ propose that the trait-relevance of a situation is critical for the observation of trait-related differences in behavior^32,34,35^. Recent empirical evidence extends these theories to the neural level, suggesting that the trait-relevance of tasks can also facilitate the detection of brain-trait relationships^36–38^, potentially by exposing the trait-related neural features, similar to a ‘stress-test’ for the brain (stress-test argument^23,39^). Although it is well established that certain features of movie stimuli (e.g., sensory integrity; emotional or social stimulus content) can modulate the relationship between inter-subject synchrony in brain activity during naturalistic stimuli and trait-like factors^30,40–45^, it remains unknown whether the *trait-relevance of movie scenes*, particularly in relation to neuroticism, can similarly influence the detectability of brain-trait representational similarity.

We applied IS-RSA to a large sample from the Human Connectome Project^46^ (HCP; *N* = 174; split in *N_main_ = 89; N_replication_ = 85;* Main Study) to determine whether participants’ similarity in neuroticism is reflected in the similarity of their brain activity during movie watching. Further, we evaluated which model of trait similarity best represents the brain-trait relationship. Given that neural characteristics of neuroticism might be more pronounced in trait-relevant contexts, we also investigated whether the brain-trait representational similarity - at the whole-brain-, network-, and region-specific levels - is influenced by participant-rated trait-relevance of movie scenes (independent *N* = 86; Pilot Study).

## Results

### Pilot Study identifies neuroticism-relevant movie scenes

To evaluate the trait-relevance of movie scenes, an online Pilot Study was conducted prior to the Main Study. Specifically, 86 participants rated ∼30-second scenes from the four movies presented to the Main Study participants in the fMRI scanner (each approximately 15 minutes long; Fig. 1a, Supplementary Tables S1-S2) regarding their potential to evoke neuroticism-related emotions.

**Fig. 1.**
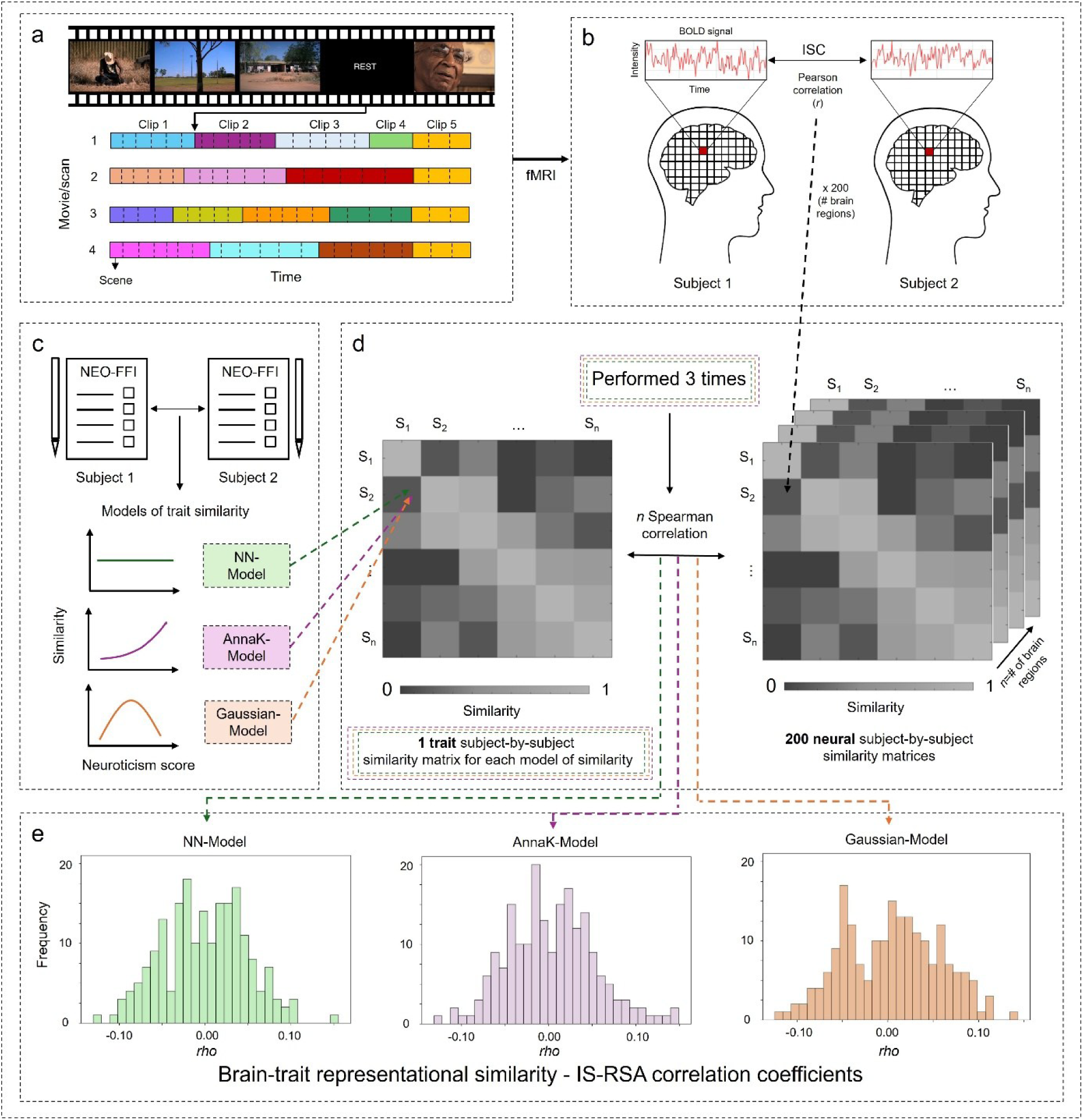
Schematic overview of study procedure. **(a)** Individual brain activity was recorded while participants watched four movies, each containing four to five clips which were separated by 20-second rest periods (Supplementary Table S1). To assess trait-relevance of movie stimuli in the Pilot Study, clips were further divided into scenes with durations of approximately 30 seconds (Supplementary Table S2). **(b)** Inter-subject similarity in brain activity was assessed via Pearson correlations between brain region-specific activity time courses recorded during movie watching (Inter-subject correlation^20,25^; Figure panel adapted from Hasson Lab^70^) **(c)** Neuroticism was measured with the NEO-FFI and inter-subject similarity in neuroticism was assessed with three distinct models: Nearest Neighbors Model, Anna Karenina Model, and Gaussian Model. **(d)** Inter-subject representational similarity analysis (IS-RSA^22^) was applied to test whether the inter-subject similarity in neuroticism is represented in the inter-subject similarity (i.e., synchrony) of brain activity. Specifically, the *trait* subject-by-subject similarity matrix for each of the three models of trait similarity was compared to each region-specific *neural* subject-by-subject similarity matrix (*N* = 200) via Spearman correlation. **(e)** This yielded a set of 200 region-specific IS-RSA correlation coefficients for each model of trait similarity (note that depicted distributions were randomly generated only for visualization purposes). BOLD = blood oxygen level dependent; ISC = inter-subject correlation; IS-RSA = inter-subject representational similarity analysis.

The Pilot Study sample consisted of two groups, which were matched to the Main Study sample in terms of age and gender distribution. Group 1 (*N_group1_* = 41; 24 females; mean age = 30.9 years; age range = 23 - 36 years) rated the 44 scenes from Movies 1 and 2, while Group 2 (*N_group2_* = 45; 25 females; mean age = 30.4 years; age range = 22 - 36 years) rated the 44 scenes from Movies 3 and 4. Neuroticism scores, assessed with the NEO-Five-Factor Inventory (NEO-FFI)^47,48^, ranged from 2 - 39 (*M* = 21.7; *SD* = 9.2) in Group 1 and from 6 - 45 (*M* = 23.9; *SD* = 8.4) in Group 2 and were approximately normally distributed in both groups (Shapiro-Wilk test: Group 1 (*W*(41) = 0.978, *p* = .591), Group 2 (*W*(45) = 0.985, *p* = .811; frequency distribution in Supplementary Fig. S1a-b).

For each participant, a single score per scene was computed based on their scene ratings (responses to six questions presented after each scene; Supplementary Table S3; Supplementary Fig. S2), with higher scene scores indicating greater trait-relevance.

Distributions of scene scores are illustrated in Supplementary Fig. S3. Within both conditions (i.e., scenes rated by Group 1 or Group 2, respectively), all scenes classified as trait-relevant (top quartile of mean scene scores) had significantly higher mean scene scores than all scenes classified as trait-irrelevant (bottom quartile of mean scene scores; one-way repeated-measures ANOVA followed by Holm-adjusted pairwise comparisons; all *p* < 0.05, Supplementary Fig. S4) and participants were highly attentive, thus providing an important prerequisite for the Main Study: 51.2% (*N* = 44) strongly agreed and 43.0% (*N* = 37) agreed with the statement “*I was able to stay focused and pay close attention […]*”; while 4.7% (*N* = 4) were neutral, and only 1.2% (*N* = 1) disagreed.

### Main Study reveals that neuroticism-relevant movie scenes accentuate neuroticism-relevant brain activity

The Main Study assessed brain-trait representational similarity, that is whether participant’s similarity in neuroticism is reflected in the similarity of their brain activity during movie watching (Fig. 1). For this purpose, data from 174 HCP participants (104 females; mean age = 29.3 years; age range = 22 - 36 years) was used and information about the trait-relevance of movie scenes as identified in the Pilot Study was incorporated.

In the main sample (*N_main_* = 89; 53 females; mean age = 29.4 years; age range = 22 - 36 years), NEO-FFI^47,48^-assessed neuroticism scores ranged from 3 - 39 (*M* = 16.5; *SD* = 6.6) and were approximately normally distributed (Shapiro-Wilk test: *W*(89) = 0.980, *p* = 0.177; frequency distribution in Supplementary Fig. S1c).

#### Different models of inter-subject similarity in neuroticism inform distinct assumptions about the brain-trait representational similarity

Inter-subject similarity in neuroticism was assessed with three different models: The Nearest Neighbors (NN)-Model (similarity in brain activity is assumed to correspond to similarity in neuroticism scores), the Anna Karenina (AnnaK)-Model (similarity in brain activity is assumed to increase, as neuroticism scores increase), and the Gaussian-Model (similarity in brain activity is assumed to be greatest in mid-range neuroticism scorers, Fig. 1c)^22^. This yielded three *trait* subject-by-subject matrices, which provide the behavioral input data for the IS-RSA. Visual representations of the intended similarity structures are represented in Fig. 2a, where subjects are presented on the axes and sorted by neuroticism rank (from low to high; left-hand side panels).

**Fig. 2.**
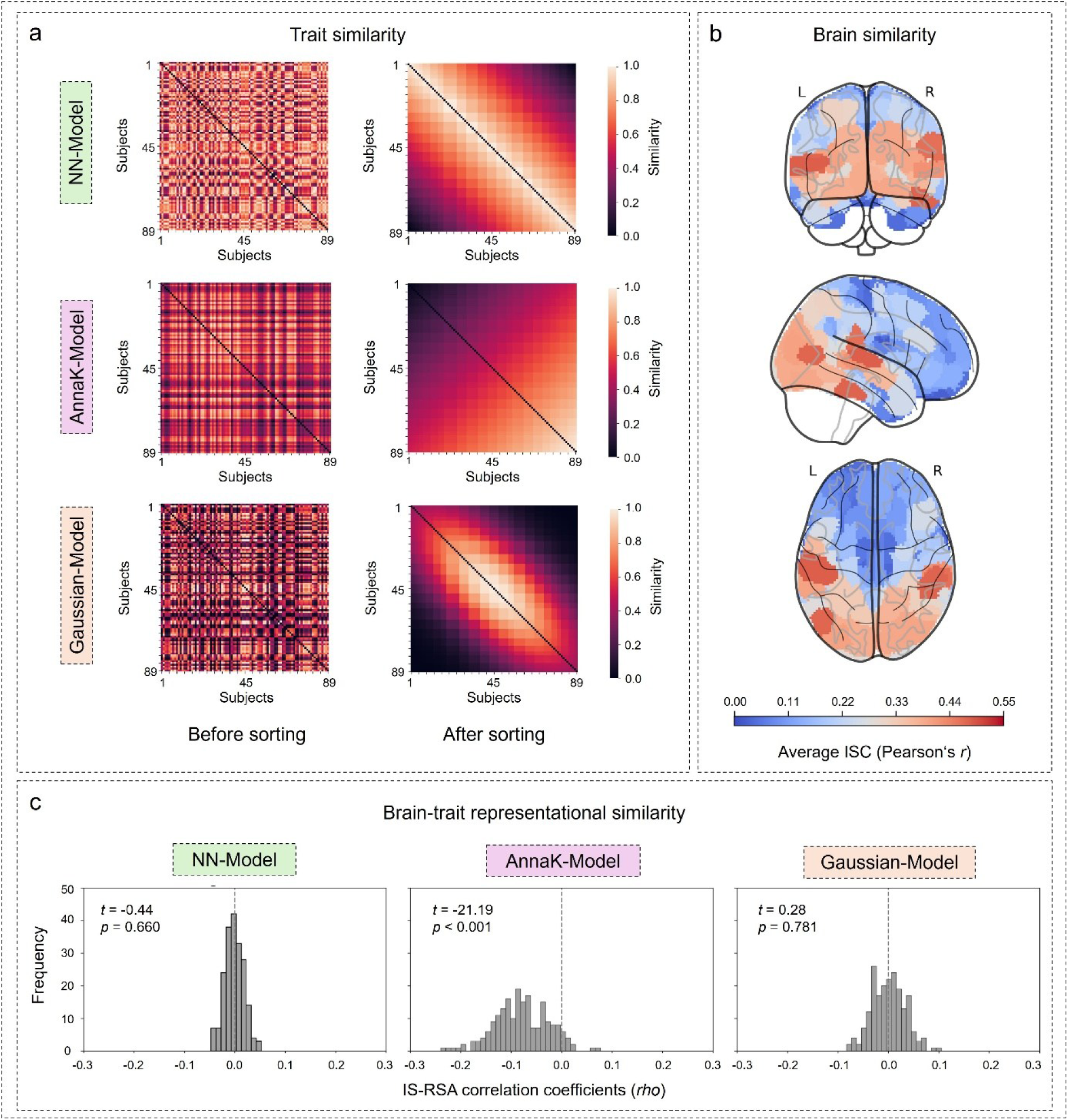
Inter-subject representational similarity analysis in the main sample (*N* = 89) **(a)** Inter-subject trait similarity in the main sample is presented for three trait similarity models implementing different assumptions about the form of brain-trait representational similarity (NN-Model, AnnaK-Model, and Gaussian-Model). In the matrix on the left, subjects are ordered as in the dataset. On the right, the same matrix is shown with subjects sorted by neuroticism rank (from low to high), highlighting the underlying inter-subject similarity structure in neuroticism. **(b)** Brain map reflecting the overall similarity of neural responses across all participant pairs. Specifically, region-specific time courses of brain activity from all scenes were correlated for each pair of participants (ISC). The resulting region-specific ISC correlation coefficients were averaged across all participant pairs to produce a general map illustrating the whole-brain pattern of inter-subject similarity in brain activity. Higher inter-subject synchrony in brain activity was observed in unimodal sensory regions, whereas multimodal association regions exhibited lower synchrony. **(c)** Frequency distribution of model-specific IS-RSA correlation coefficients computed by comparing the lower triangles of the model-specific *trait* subject-by-subject similarity matrix to each of the 200 brain region-specific *neural* subject- by-subject similarity matrices via Spearman correlation. One-sample *t*-tests were performed to test whether the distribution was significantly shifted from zero, indicating representational brain-trait similarity at the whole-brain level. ISC = inter-subject correlation; IS-RSA = inter-subject representational similarity analysis.

#### Inter-subject similarity in brain activity follows cortical functional organization

After preprocessing (see Methods), fMRI-recorded time courses of brain region-specific activity from all four movies were compiled separately for each of the 200 brain regions^49^. These 200 aggregated time series were then compared across all subjects (ISC^20,25^; Fig. 1b) to assess region-specific inter-subject similarity in brain activity. This yielded 200 *neural* subject-by-subject matrices. The group-general whole-brain pattern of similarity in brain activity was visualized by averaging region-specific ISC correlation coefficients across all participant pairs (Fig. 2b). Overall, higher inter-subject neural similarity was observed in unimodal areas engaged in audiovisual processing, whereas lower similarity was detected in multimodal association regions associated with higher-order processing.

#### Brain-trait representational similarity is best captured by the inverse AnnaK-Model

Brain-trait representational similarity was assessed by comparing model-specific *trait* subject-by-subject similarity matrices to each brain region-specific *neural* subject-by-subject similarity matrix via Spearman correlation, yielding three sets of model-specific 200 IS-RSA correlation coefficients (Fig. 1d-e). Taking brain activity from all 200 brain regions during all movie scenes (3086 frames) into account, significant brain-trait representational similarity was only detected under the AnnaK-Model, where IS-RSA correlation coefficients were significantly different from zero (*t*(199) = -21.19, *p* < 0.01; Fig. 2c). As IS-RSA values were shifted to the left (mostly negative), this indicates that the inverse of what is assumed under the AnnaK-Model applies: Low neuroticism scorers have similar brain activity, while high neuroticism scorers differ in their brain activity from each other. Put into other words: As neuroticism scores increase, the heterogeneity in brain responses increases.

For the NN-Model and the Gaussian-Model, brain-trait representational similarity was not significant (all *p* > .05; Fig. 2c) when considering all movie scenes and all brain regions. Therefore, in the following, we focused on the comparison of brain-trait representational similarity between trait-relevant and trait-irrelevant scenes under the AnnaK-Model. Results based on the NN-Model and Gaussian-Model are reported in the Supplementary Information (Supplementary Tables S4-S7).

#### Higher brain-trait representational similarity during trait-relevant scenes

To compare brain-trait representational similarity between trait-relevant and trait-irrelevant scenes, corresponding brain region-specific activity time courses were compiled across all trait-relevant scenes (843 frames) and trait-irrelevant scenes (708 frames). Based on these compiled time series, brain region-specific *neural* subject-by-subject similarity matrices were constructed via ISCs (Fig. 1d). The group-general whole-brain patterns of similarity in brain activity were again visualized by averaging region-specific ISC correlation coefficients across all participant pairs (Supplementary Fig. S5a). Overall, these patterns were highly similar between trait-relevant and trait-irrelevant scenes.

When collectively considering brain activity in all 200 brain regions (i.e., whole-brain level), significant brain-trait representational similarity (under the AnnaK-Model) was detected for both trait-relevant scenes (*t*(199) = -20.35, *p* < 0.01; Fig. 3a; Supplementary Table S4) and trait-irrelevant scenes (*t*(199) = -9.10, *p* < .001; Fig. 3a; Supplementary Table S5). However, the respective sets of IS-RSA correlation coefficients significantly differed (*t*(199) = -10.87, *p* < .001), which indicates greater brain-trait representational similarity during trait-relevant scenes compared to trait-irrelevant scenes.

**Fig. 3.**
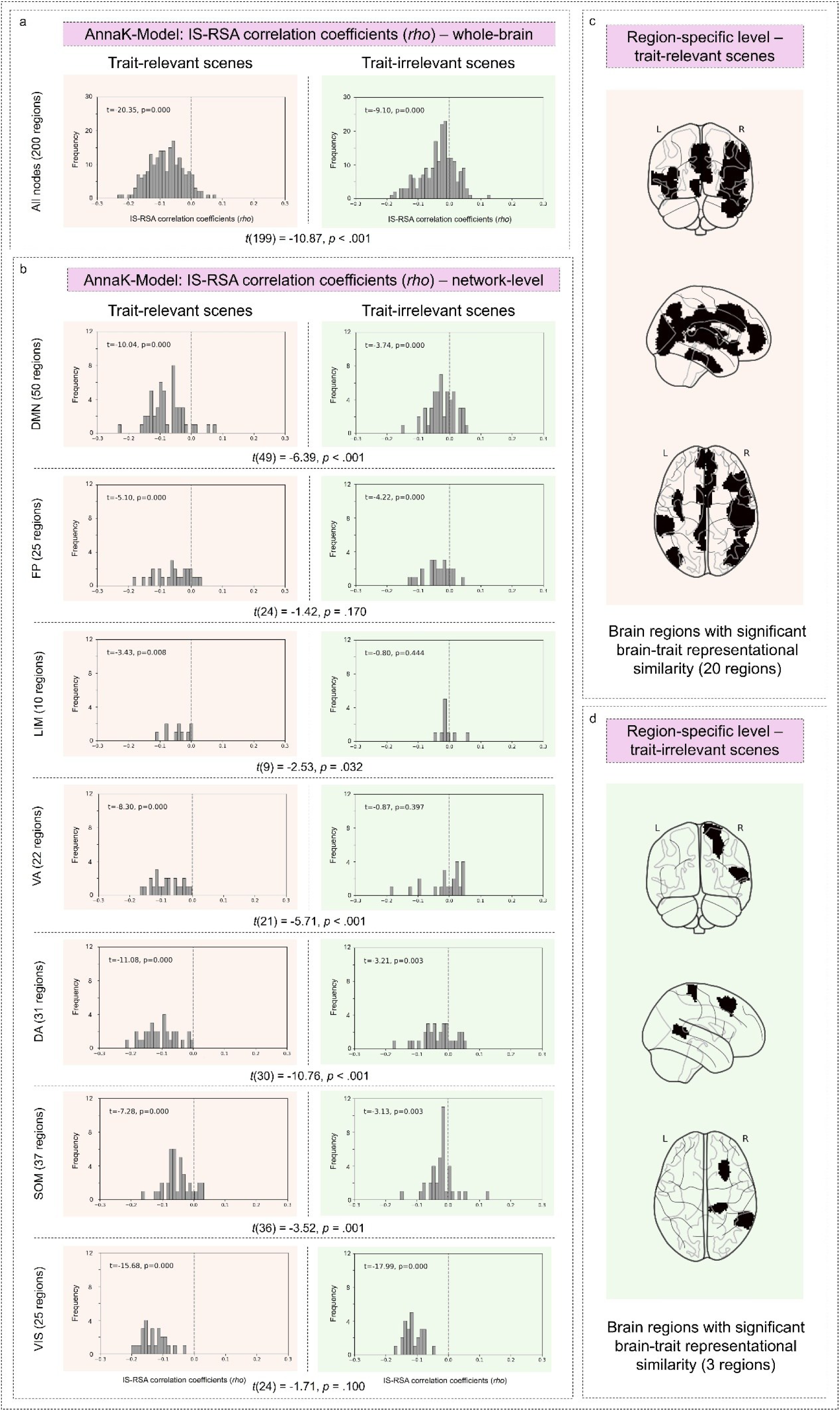
Comparison of brain-trait representational similarity under the Anna-K Model between trait-relevant and trait-irrelevant scenes. **(a)** Frequency distribution of IS-RSA correlation coefficients from the assessment of brain-trait representational similarity during trait-relevant and trait-irrelevant scenes under the AnnaK-Model when considering all 200 regions of the whole brain simultaneously. One-sample *t*-tests were performed to test whether each distribution was significantly shifted from zero, indicating representational brain-trait similarity at the whole-brain level. Further, a paired-sample *t*-test was performed with both sets of 200 IS-RSA correlation coefficients to assess whether brain-trait representational similarity significantly differed between trait-relevant and trait-irrelevant scenes. **(b)** Frequency distributions of IS-RSA correlation coefficients, split by the Yeo seven-network affiliation^50^, such that each analysis and respective panel only includes the coefficients corresponding to brain regions within the respective network. One-sample *t*-tests were performed to assess whether each distribution is significantly shifted from zero, indicating representational brain-trait similarity at the network-level. Further, paired-sample *t*-tests on both respective sets of network-specific IS-RSA correlation coefficients were conducted to test for significant differences in brain-trait representational similarity between trait-relevant and trait-irrelevant scenes. **(c)** Brain regions with significant brain-trait representational similarity during trait-relevant scenes were identified via non-parametric permutation testing (Mantel-test; uncorrected threshold of *p* < 0.05). Corresponding IS-RSA correlation coefficients and *p*-values are shown in Supplementary Table S6. No brain region remained significant after FDR correction for multiple comparisons. **(d)** Single brain regions with significant brain-trait representational similarity during trait-irrelevant movie scenes (Mantel-test; uncorrected threshold of *p* < 0.05). Corresponding IS-RSA correlation coefficients and *p*-values are shown in Supplementary Table S7. Again, no brain region remained significant after FDR correction. IS-RSA = inter-subject representational similarity analysis; DMN = default mode network; FP = frontoparietal control network; LIM = limbic network; VA = ventral attention/salience network; DA = dorsal attention network; SOM = somatomotor network; VIS = visual network.

At the level of the seven functional Yeo networks^50^ (Supplementary Fig. S6), significant brain-trait representational similarity during trait-relevant movie scenes was observed in the a) default mode network (DMN; *t*(49) = -10.04, *p* < 0.001), b) frontoparietal control network (FP; *t*(24) = -5.10, *p* < 0.001), c) ventral attention/salience network (VA; *t*(21) = -8.30, *p* < 0.001), d) dorsal attention network (DA; *t*(30) = -11.08, *p* < 0.001), e) somatomotor network (SOM; *t*(36) = -7.28, *p* < 0.001), and f) the visual network (VIS; *t*(24) = -15.68, *p* < 0.001) (Fig. 3b; Supplementary Table S4). During trait-irrelevant scenes, brain-trait representational similarity was significant in the a) DMN (*t*(49) = -3.74, *p* < 0.001), b) FP (*t*(24) = -4.22, *p* < 0.001), c) DA (*t*(30) = -3.21, *p* = 0.003), d) SOM (*t*(36) = -3.13, *p* = 0.003), and e) the VIS (*t*(24) = -17.99, *p* < .001) (Fig. 3b; Supplementary Table S5). Brain-trait representational similarity was significantly greater during trait-relevant compared to trait-irrelevant scenes in the a) DMN (*t*(49) = -6.39, *p* < 0.001), b) VA (*t*(21) = -5.71, *p* < 0.001), c) DA (*t*(30) = -10.76, *p* < 0.001), and d) SOM (*t*(36) = -3.52, *p* < 0.001) (Fig. 3b). This corroborates the whole-brain findings. No significant difference was observed in the a) FP (*t*(24) = -1.42, *p* = .170), b) LIM (*t*(10) = -2.53, *p* = .032), and c) VIS (*t*(24) = -1.71, *p* = .10) (Fig. 3b).

At the level of the 200 separate brain-regions, significant brain-trait representational similarity was detected in 20 regions during trait-relevant scenes and in three regions during trait-irrelevant scenes (non-parametric Mantel test^51^; *p* < 0.05, Fig. 3c-d; Supplementary Tables S6-S7). However, none of these regions remained significant after FDR correction for multiple comparison (significance level of *p* < .05).

### Replication

To assess robustness and generalizability of study findings, all analyses were repeated in a replication sample (*N_replication_* = 85; 51 females; mean age = 29.2 years; age range = 22 - 36 years). Neuroticism scores ranged from 2 - 31 (*M* = 16.2; *SD* = 6.0) and were approximately normally distributed (Shapiro-Wilk test: *W*(85) = 0.987, *p* = 0.565; frequency distribution in Supplementary Fig. S1d).

Inter-subject similarity in neuroticism was similarly operationalized for the three models as in the main sample and again showed the intended similarity structure after sorting (Supplementary Fig. S7a). The group-general whole-brain pattern of inter-subject similarity in brain activity across all movie scenes in the replication sample was also highly similar to the main sample (Supplementary Fig. S7b).

When assessing brain activity from all 200 brain regions simultaneously during all movie scenes, significant brain-trait representational similarity was only detected under the AnnaK-Model (*t*(199) = -17.99, *p* < 0.001; Supplementary Table S8; Supplementary Fig. S7c), thereby corroborating the main analyses.

The group-general whole-brain patterns of similarity in brain activity detected during all trait-relevant and all trait-irrelevant movie scenes in the replication sample were also highly similar to the main sample (Supplementary Fig. S7b).

For trait-relevant scenes, significant brain-trait representational similarity was replicated at the whole brain level, in the DMN, FP, VA, DA, SOM, and in the VIS (Supplementary Table S4; Supplementary Fig. S8a-b). For trait-irrelevant scenes, brain-trait representational similarity was replicated at the whole-brain level, in the DMN, FP, DA, SOM, and in the VIS (Supplementary Table S5; Supplementary Fig. S8a-b). Significantly greater brain-trait representational similarity during trait-relevant compared to trait-irrelevant scenes could be replicated at the whole-brain level, in the DMN, and in the SOM (Supplementary Fig. S8a-b). At the region-specific level, none of the uncorrected results from the main sample were replicated, apart from one single brain region (RH_SalVentAttnB_PFCv_2) which showed significant brain-trait representational similarity during trait-relevant movie scenes (Supplementary Tables S6-S7; Supplementary Fig. S8c-d).

### Post-hoc analyses

To further assess the robustness of region-specific brain-trait representational similarity, model- and brain region-specific IS-RSA correlation coefficients derived from the main analysis were plotted against those from the replication analysis and compared using Pearson correlations (Supplementary Fig. S9). Significant positive correlations were observed for values from all scenes (*r* = 0.34; *p* < 0.001), trait-relevant scenes (*r* = 0.34; *p* < 0.001), and trait-irrelevant scenes (*r* = 0.27; *p* < 0.001).

For completeness, results from analyzing network- and region-specific brain-trait representational similarity under all three trait similarity models during all scenes are reported in Supplementary Tables S8-S9.

## Discussion

By applying IS-RSA to 174 participants from the HCP (Main Study; *N*_main_ *= 89; N*_replication_ *= 85)*, this study examined a) whether individuals’ similarity in neuroticism is reflected in the similarity of their brain activity during movie watching, and b) whether this brain-trait representational similarity depends on the trait-relevance of movie scenes, while trait-relevance was rated in an independent Pilot Study (*N* = 86). Overall, inter-subject similarity in brain activity was higher in unimodal sensory regions and lower in multimodal association regions. Brain-trait representational similarity was best captured by the inverse AnnaK-Model, implying that heterogeneity in brain activity increases with higher neuroticism scores. Supporting our preregistered hypotheses, brain-trait representational similarity was significantly greater during trait-relevant compared to trait-irrelevant movie scenes. These findings substantiate contemporary personality theories, which propose that individual differences in personality traits at the behavioral level are better observable in trait-relevant situations, from a neural perspective. We conclude by underscoring the importance of stimulus selection when investigating brain-trait relationships and by highlighting the potential of naturalistic neuroimaging for studying mechanisms underlying mental disorders, specifically in patient populations.

The successful identification of scenes as relevant or irrelevant to the trait neuroticism in our Pilot Study aligns with prior research demonstrating that emotionally salient movie clips elicit consistent interpretations and shared affective experiences across individuals^52–54^. Although this provides a reliable foundation for investigating how exposure to trait-relevant stimuli influences brain-trait representational similarity in the Main Study, the specific visual and auditory properties of the movie (i.e., sensory intensity, social content, number of spoken words) that contribute to a movie’s ability to evoke neuroticism- or other trait-relevant emotions remain to be explored in future studies.

Overall, the similarity in movie-evoked brain activity between individuals was higher in unimodal sensory regions and lower in multimodal association regions. Aligning with previous research^20,22,42^, this reflects that heterogeneity in movie-evoked brain activity is greater in regions at the top of the principal gradient of macroscale cortical organization^55^ and is compatible with their suggested critical role for higher-order functions: Unimodal sensory regions involved in audiovisual information processing are thought to closely track the dynamic features and temporal structure of the movie stimuli, producing highly similar response patterns across individuals who are simultaneously exposed to the same sensory input^20,42,56^. In contrast, as regions move up in the cortical processing hierarchy^55^ and take on higher-order functions such as emotion processing, their neural signal contains less of this consistent stimulus-evoked response and is increasingly dominated by individual, potentially trait-related patterns. This would then lead to decreased inter-subject similarity in brain acivity^22^. Replicating Li et al.^42^, the group-average pattern of whole-brain inter-subject similarity in brain activity was relatively independent of scene content. However, this rather stable group-general global pattern does not preclude the presence of small but trait-related differences in inter-subject similarity in brain activity, which are potentially embedded in the fine-grained variability of context-dependent neural similarity among participant pairs.

The inverse AnnaK-Model best captured the representational similarity between neuroticism and brain activity. This implies that movie watching-related brain activity becomes more heterogeneous as neuroticism increases. From a mechanistic perspective, this heterogeneity may stem from neurotic individuals’ tendency to focus on negative emotions. Emotionally charged scenes could trigger increased rumination and worry, driving neurotic individuals’ attention inward and activating memory content related to personal experiences (autobiographical memories). This, of course, differs between individuals and could thus explain the increased heterogeneity in brain activity in persons with high neuroticism. Moreover, this sustained engagement with internal thoughts consumes cognitive resources, potentially impairing attention to the stimulus^57,58^. As a result, the brain activity of individuals high in neuroticism may diverge from both low neuroticism scorers, who probably process the stimuli more uniformly and therefore show relatively similar brain activity relative to each other, and from other high scorers, whose attention is directed differently.

With that, our findings also extend previous research on inter-subject similarity in brain activity in clinical and subclinical mental health conditions, suggesting that deviations from the norm are associated with an enhanced risk for malfunctioning: During movie watching, inter-subject similarity in brain activity was not only shown to be diminished in patients with melancholic depression^58^, and first order psychosis^59^, but also in children with higher social anxiety symptoms^40^ and adolescents with heightened depressive symptoms^30^. The concordance between our findings regarding neuroticism and those from clinical populations not only substantiates the well-established link between heightened neuroticism and mental disorders, but also informs a dimensional understanding of psychopathology^60^, where psychopathology is considered as the extreme part of the normal distribution of a trait. A deviation from ‘typical‘ brain activity during naturalistic stimulation compared to healthy populations, as demonstrated in our study, may emerge even before the onset of a clinical disorder, thus serving a promising early neural indicator that could be used to identify high-risk populations and to mitigate disease onset via personalized preventions.

Contrary to our findings, electroencephalography (EEG) research has revealed a positive linear association between inter-subject neural similarity and inter-subject neuroticism similarity^31^, suggesting that brain-trait representational similarity may be sensitive to the spatial and temporal resolution of the measurement technique. Speculatively, individuals similar in neuroticism may process emotionally salient stimuli temporally similar (better captured by EEG), while spatial activation patterns might differ (better captured by fMRI). This method-dependency in brain-trait representational similarity presents an interesting question for future research.

Lastly, the lack of significant region-specific associations in the presence of whole-brain and network-level effects aligns with research on intelligence as another complex human trait^37,61^, supporting that personality differences are represented in widespread functional networks rather than individual brain regions^15^.

Current personality theories emphasize interactions between traits and situations^32,33^. By revealing greater representational similarity between neuroticism scores and brain activity during trait-relevant compared to trait-irrelevant scenes, we support the extension of these theories to the neural level. Specifically, we propose that neuroticism-relevant movie scenes may act as situational triggers engaging neural processes inherent to neuroticism: The engagement of neuroticism-relevant neural features increases their detectability, as well as the detectability of related individual differences in these features, similar to a neural stress-test^23,39^. This pronounced detectability of each individuals’ neural neuroticism signature could ultimately amplify brain-trait representational similarity. Support for this interpretation also comes from previous neuroimaging research focused on intelligence and other personality traits, showing that brain-trait associations become more detectable when neural activity is constrained by trait-relevant task paradigms^36–38^. Collectively, these results also extend previous empirical evidence for recent theoretical work proposing trait-relevant task selection as promising means to increase power in neuroimaging research on individual differences^62^. From a broader perspective, the here applied framework - IS-RSA - offers a straightforward means for testing hypotheses derived from personality theories.

In general, naturalistic stimuli are particularly well-suited for patient populations^19^: They promote engagement and reduce participants’ awareness of the scanner environment, thereby enhancing subject compliance and improving data quality compared to resting-state paradigms. Additionally, they require no prior training, reduce task performance-related pressure, and allow for flexible adaptation of stimulus content and intensity. Our work further substantiates the potential of naturalistic neuroimaging not only for investigating the neurobiological underpinnings of personality traits, but - given the link between neuroticism and psychological disorders - also for studying mental health conditions. By selecting stimuli tailored to specific clinical traits or maladaptive behaviors, relevant systems could be effectively engaged, thus providing more specific insights into the underlying brain dysfunction. This could ultimately assist in diagnosis or monitoring of treatment^16,58^.

## Limitations

This study has limitations. Most importantly, we could not perform replication analyses in an independent data set, as no suitable open-access data set is currently available. Second, while we consider it a strength that trait-relevance of movie scenes was rated by an independent sample, it is uncertain whether movies might have induced the same experiences in the main sample during fMRI scanning. Further, trait-relevance was evaluated post hoc, after stimulus presentation, thus inducing the potential of memory-bias, and our study was limited by the specific movies selected by the Human Connectome Project. Future studies could benefit from explicitly designing stimuli, such as movies, tailored to the trait of interest and by performing rating procedures in the neuroimaging sample and an independent sample live and post-hoc for validation. Third, no direct measure of participant attention was included in the Main Study (e.g., eye-tracking or post-scanning questionnaires), leaving the possibility that differences in attentional focus during neuroimaging might have influenced the results. Finally, we could not control for potential differences in the familiarity with movie content, which might have affected individuals’ attention and their perception in both the Main Study and the Pilot Study.

## Conclusion

Our findings revealed that higher levels of neuroticism were associated with greater heterogeneity in brain activity during naturalistic stimuli, with this effect being strongest when stimuli were trait-relevant. This increased neural heterogeneity may be driven by reactivation of idiosyncratic memories, attention deficits, and emotion dysregulation. First, our results mirror what has been observed in patient populations, where deviations from normative neural activity patterns are commonly dysfunctional, thus supporting dimensional conceptualizations of psychopathology that position high neuroticism as risk factor for mental illness. Second, the amplification of brain-trait representational similarity during trait-relevant movie scenes corroborates contemporary personality theories that emphasize the essential role of situational context in observing personality differences - at the neural level. More generally, our results highlight the potential of naturalistic neuroimaging to advance our understanding of the neural basis underlying personality traits and psychopathology, while underscoring the importance of stimulus selection in the investigation of brain-behavior associations.

## Methods

### Preregistration

This study was preregistered on the Open Science Framework: https://osf.io/j7vbu. It consists of an online Pilot Study, rating movie scenes for their potential to evoke neuroticism-related emotions, and a Main Study, assessing brain-trait representational similarity, while incorporating the trait-relevance as defined in the Pilot Study.

### Movie stimuli

The same set of four movies was shown in both the Pilot Study and the Main Study. Each movie was approximately 15 minutes long and consisted of four to five independent clips ranging from 1:04 - 4:18 minutes (Fig. 1a). Movies 1 and 3 contained clips from independent films, including both fiction and documentary, while Movies 2 and 4 featured clips from Hollywood productions (Supplementary Table S1; Cutting et al., 2012). In the Pilot Study, the clips were further segmented into 88 scenes of ∼30 seconds length (*M* = 36.1 s; *SD* = 10.33 s; range = 17 - 64 s; Fig. 1a). Scene durations varied slightly to preserve the narrative flow, and minor adjustments (i.e., trimming frames) were made to ensure smooth transitions between scenes (Supplementary Table S2). In the Main Study, the four movies were presented inside the fMRI scanner, with 20-second rest periods between clips, during which the word ‘REST’ was displayed on the screen. Additional 20-second rest periods were included at the beginning and end of each movie.

### Pilot Study: Identifying trait-relevant movie scenes

#### Participants

Participants were recruited from the general population in the United States via an online platform (www.clickworker.com). Following Gruskin et al.^30^, where 20 adults rated the emotional valence of a 10-minute clip, we aimed to recruit 40 participants for the scene rating procedure, with the goal of matching the age range (22 - 36 years) and gender distribution (60% female, 40% male) of the Main Study. To maintain participants’ attention, we limited the session length to approximately one hour, by dividing the movies’ scenes between two groups: One group (*N* ≈ 40) rated scenes from Movies 1 and 2, while the other group (*N* ≈ 40) rated scenes from Movies 3 and 4.

Eligibility criteria included normal or corrected-to-normal vision and hearing, as well as proficiency in English. Participants confirmed that they were neither diagnosed with a psychological or psychiatric condition, nor under the influence of alcohol, illicit drugs, or medications that could affect scene perception. The final sample for Group 1 comprised 41 participants (24 females; mean age = 30.9 years; age range = 23 - 36 years). After excluding one participant due to uniform ratings across all scenes, most likely indicating a lack of engagement with the experimental instructions, the final sample for Group 2 comprised 45 participants (25 females; mean age = 30.4 years; age range = 22 - 36 years).

#### Procedure

Participants were instructed to complete the online study on a computer screen and to ensure a quiet location with access to headphones or high-quality speakers. First, participants were informed about the study’s purpose and procedure, provided informed consent, and reported demographic information including age, gender, highest level of education, years of education, and occupation. Second, they completed the NEO-Five-Factor Inventory (NEO-FFI), a 60-item self-report questionnaire based on the Costa and McCrae Five-Factor Inventory^47,48^. Particularly, the NEO-FFI contains 12 items to assess each Big Five personality trait, including neuroticism. Participants rated their agreement with statements reflecting their self-perception (e.g., “I am not easily disturbed”) on a five-point Likert scale ranging from 0 (strongly disagree) to 4 (strongly agree).

Before beginning the actual scene ratings (44 per condition), participants viewed a single test scene to confirm well-functioning video and audio setup. The rating process was structured as follows: Each scene, presented in consecutive order, was followed by six randomly ordered statements designed to assess its perceived relevance to neuroticism (Supplementary Table S3). These statements covered the six facets of neuroticism (anxiety, angry hostility, self-consciousness, depression, impulsiveness, and vulnerability) and participants rated their agreement with each statement on a five-point Likert scale ranging from 0 (strongly disagree) to 4 (strongly agree). The facet-specific statements were developed by the authors for the present study. They were created based on the NEO-FFI and state neuroticism items introduced by Beckman et al.^64^, but were slightly adapted to reflect the atmosphere and emotions conveyed in the scenes. To minimize response bias and to foster an engaging and comfortable atmosphere, half of the statements were phrased in the opposite direction, capturing both the perception of negative neuroticism-related emotions and their positive counterparts. Prior to the ratings, each statement was briefly explained to reduce ambiguity in interpretation.

Upon completing all scene ratings, participants’ attentiveness throughout the ratings was assessed with the statement “*I was able to stay focused and pay close attention to the movie scenes throughout the entire experiment”,* on a five-point Likert scale ranging from 0 (strongly disagree) to 4 (strongly agree). An overview of the Pilot Study is presented in Supplementary Fig. S2.

The online experiment was implemented using PsychoPy and Pavlovia, and participants were compensated with 10 Euros upon completion.

#### Identification of trait-relevant scenes

Per participant, a total score for each scene was computed by summing the responses to the six corresponding statements. Reverse-coded items were re-coded in advance (e.g., 0 becomes 4, 1 becomes 3, and 2 remains 2). To account for individual response tendencies, the resulting scene scores were then rescaled to a range of 1-10 within each participant (i.e., across all 44 scenes). These rescaled scores were then averaged across participants for each scene. Scenes falling into the top quartile (11 scenes per condition) were classified as trait-relevant, while those falling into the bottom quartile were classified as trait-irrelevant. Per condition, a one-way repeated-measures ANOVA followed by Holm-adjusted post-hoc pairwise comparisons (significance threshold *p* < 0.05) was performed to test whether trait-relevant and trait-irrelevant scenes differed significantly in participant ratings (i.e., scaled scene scores).

### Main Study: Assessing brain-trait representational similarity

#### Participants

The Human Connectome Project (HCP) dataset with movie-watching 7T fMRI data from 184 adult participants was used^46^. We excluded participants with incomplete neuroimaging data from any of the movies or excessive head motion (mean framewise displacement > 0.55 mm)^65^. Due to family relations, we divided the dataset into two approximately equally sized cohorts of unrelated participants^22^. Because one family contained four members, two of them were randomly excluded to ensure that cohorts only contain unrelated subjects. The final sample consisted of 174 participants (104 females; mean age = 29.3 years; age range = 22 - 36 years) and was split into a main sample (*N_main_* = 89; 53 females; mean age = 29.4 years; age range = 22 - 36 years) and a replication sample (*N_replication_* = 85; 51 females; mean age = 29.2 years; age range = 22 - 36 years). Neuroticism was assessed with the NEO-FFI^47,48^ outside the scanner.

#### FMRI data acquisition and preprocessing

To estimate brain region-specific activity time courses, the minimally preprocessed high-resolution movie fMRI data was used^46,66^. FMRI scans were collected on a 7T Siemens Magnetom scanner with a 32-channel head coil, while subjects watched four movies with audio (see **Movie stimuli**). Movies 1 and 2 were shown in a first session, while Movies 3 and 4 followed in a second session on a different day. All fMRI data were acquired with a gradient-echo planar imaging sequence (repetition time [TR] = 1000 ms; echo time [TE] = 22.2 ms; flip angle = 45°; 1.6 mm isotropic voxel resolution; multi-band factor = 5). A comprehensive description of all neuroimaging parameters can be found in the HCP’s online documentation (https://www.humanconnectome.org/study/hcp-young-adult/document/1200-subjects-data-release), and preprocessing steps are detailed in Glasser et al.^66^. In addition to the ICA-FIX denoising implemented in the HCP pipeline, mean white matter and cerebrospinal fluid traces were regressed out of vertex-wise time series data. Afterwards, brain region-specific activity time courses were extracted using the functional Yan 200 parcellation^49^, an updated version of the Schaefer parcellation^67^. We opted for the parcellation into 200 brain regions (nodes), as parcellations in the 200-300 node range have been shown to provide a good balance between spatial specificity and robustness, avoiding increased registration errors and partial voluming effects associated with smaller brain regions in higher resolution parcellations^68^.

For each clip, the first 10 frames (i.e., seconds) of brain region-specific activity time courses were excluded to mitigate the impact of large BOLD signal changes at clip onset after each rest period^23^. To account for hemodynamic delay, the 5 frames immediately after clip offset were included. The remainder of brain region-specific activity time courses recorded during the 20-second rest period between each pair of clips was discarded.

#### Inter-subject representational similarity analysis (IS-RSA)

IS-RSA^22^ assesses brain-trait representational similarity by relating inter-subject trait similarity to inter-subject similarity in region-specific brain activity.

#### Inter-subject similarity in neuroticism

Inter-subject similarity in neuroticism was assessed with three models, each implying and therefore allowing to test distinct assumptions about the form of brain-trait representational similarity^22^ (Fig. 1c):

#### Nearest Neighbors (NN)-Model

The NN-Model assumes that individuals with similar neuroticism scores also exhibit more similar brain activity, irrespective of whether they score high or low on neuroticism. In this case, inter-subject trait similarity is operationalized as the inverse of the Euclidean distance between neuroticism rank scores.

#### Anna Karenina (AnnaK)-Model

The AnnaK-Model assumes that individuals with high neuroticism scores are more similar in brain activity to each other, while low scorers are less similar in their brain activity to each other. Here, inter-subject trait similarity is operationalized as the average of both neuroticism rank scores divided by the number of subjects.

#### Gaussian-Model

The Gaussian-Model assumes that individuals with mid-range neuroticism scores are more similar in their brain activity to each other, while individuals at the extreme ranges of the distribution are less similar in their brain activity to each other. Here, inter-subject trait similarity (i.e., pairwise similarity *S_i,j_* between participant *i* and *j*) was operationalized using a Gaussian similarity kernel applied to normalized neuroticism rank scores:

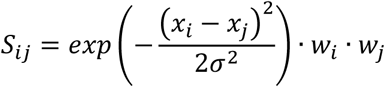

Where *x_i_* and *x_j_* are the normalized neuroticism rank scores for participant *i* and *j*, *σ* is the bandwidth of the Gaussian kernel (here, *σ* = 0.3), and *w_k_* = *exp*(−*λ* (*x_k_* − 0.5)^2^) is a middle-weighting factor for participant *k*, with *λ* = 2.

We constructed one *trait* subject-by-subject similarity matrix per model, where individual participants were placed along the axes and matrix entries depicted trait similarity between each pair of participants (Fig. 1d). Note that by assessing the effect direction (i.e., positive or negative IS-RSA correlation coefficients), IS-RSA allows to simultaneously test the original model assumption as well as its inverse: The inverse NN-Model assumes that individuals with similar neuroticism scores exhibit less similar brain activity, the inverse AnnaK-Model assumes that low neuroticism scorers are alike in their brain activity, while high scorers are less similar in their brain activity, and the inverse Gaussian-Model assumes that mid-range neuroticism scores are less similar in their brain activity, while individuals at the extreme ranges of the distribution are more similar in their brain activity.

#### Inter-subject similarity in brain activity

Inter-subject similarity in brain activity was assessed with Pearson correlations between brain region-specific activity time courses during movie watching (Inter-subject correlations, ISCs^20,25^; Fig. 1b). To this end, preprocessed and trimmed brain region-specific activity time courses from all four movies were first compiled for each participant to yield one long activity time course. The ISCs yielded one *neural* subject-by-subject similarity matrix for each of the 200 investigated brain regions, where participants were again represented along the axes, and matrix entries depicted each pair of participants’ similarity with respect to their brain region-specific activity time course (Fig. 1c). The group-general whole-brain pattern of similarity in brain activity was visualized by averaging brain region-specific similarity in neural activity across all participant pairs.

#### Assessment of brain-trait representational similarity

Per trait similarity model, brain-trait representational similarity was assessed by computing Spearman correlations between the vectorized lower triangles of the model-specific *trait* subject-by-subject similarity matrix and those of the 200 region-specific *neura*l subject-by-subject similarity matrices. This yielded 200 region-specific IS-RSA correlation coefficients for each model of trait similarity (Fig. 1e).

To statistically evaluate whether significant representational similarity exists between neuroticism and brain activity during movie watching at the whole-brain level, one-sample two-sided *t*-tests were performed on model-specific IS-RSA correlation coefficients, with Bonferroni correction applied for nine comparisons (adjusted significance threshold *p* < .006).

#### Trait-relevant vs. trait-irrelevant scenes

To compare brain-trait representational similarity between trait-relevant and trait-irrelevant scenes, we first compiled the preprocessed and trimmed region-specific activity time courses for each participant - separately for trait-relevant and trait-irrelevant scenes (defined in Pilot Study). Within this step, we also accounted for five seconds of hemodynamic delay when translating between movie scene timestamps and frames of brain region-specific activity time courses. For trait-relevant and trait-irrelevant scenes separately, the group-average whole-brain pattern of similarity in brain activity was visualized and brain-trait representational similarity was assessed as previously described, yielding 200 brain region-specific IS-RSA correlations coefficients for each trait similarity model.

#### Whole-brain level

For comparison at the whole-brain level, paired-sample two-sided *t*-tests between model-specific sets of IS-RSA correlation coefficients from trait-relevant and trait-irrelevant scenes were performed (Bonferroni-corrected threshold for eight comparisons: *p* < .006)

#### Network-specific level

For comparison at the level of functional networks, the 200 brain region-specific IS-RSA correlation coefficients were assigned to one of seven Yeo networks^50^ (Supplementary Fig. S6): DMN (50 regions), FP (25 regions), limbic network (LIM; 10 regions), VA (22 regions), DA (31 regions), SOM (37 regions), or VIS (25 regions). Model- and network-specific sets of IS-RSA correlation coefficients from trait-relevant and trait-irrelevant scenes were compared with paired-sample two-sided *t*-tests (Bonferroni-corrected threshold for eight comparisons: *p* < .006).

#### Brain region-specific level

A non-parametric Mantel test^51^ was applied to assess significance of brain-trait representational similarity at the brain region-specific level. This is necessary because values in each *subject-by-subject* similarity matrix reflect a pair of participants and are therefore not independent. Participant labels of the *trait* subject-by-subject similarity matrix were permuted 10,000 times to generate a null distribution of IS-RSA correlation coefficients. Per brain region, the observed true IS-RSA correlation coefficient was compared to this null distribution to determine a *p*-value. These *p*-values were then FDR corrected^69^, with a significance level of *p* < 0.05. For comparison at the brain region-specific level, the numbers of significant model- and brain region-specific IS-RSA correlation coefficients from trait-relevant and trait-irrelevant scenes were descriptively contrasted.

#### Replication

To evaluate robustness and generalizability of findings, all analyses were first conducted on the main sample (*N_main_* = 89) and then repeated in the lockbox sample (*N_replication_* = 85).

### Inclusion and ethics statement

For the Main Study using data from the Human Connectome Project, the Washington University Institutional Review Board^46^ approved all procedures and all participants provided informed written consent in accordance with the principles of the declaration of Helsinki. For the Pilot Study, procedures were also in consent with the Declaration of Helsinki and approved by the ethics committee of the Department of Psychology at the University of Würzburg (GZEK 2025-05; 28.02.2025). All participants provided informed written consent.

### Data availability statement

Data from the Main Study (Human Connectome Project) can be downloaded under https://www.humanconnectome.org/study/hcp-young-adult. Data from the Pilot Study was made available on the Open Science Framework: https://osf.io/j6ry5/files.

### Code availability statement

The code for all analyses, including preprocessing, is available on Github: fMRI preprocessing: https://github.com/faskowit/hcp7t_mov_proc/tree/main; Pilot Study and Main Study analyses: https://github.com/johannaleapopp/Neural_Code_of_Neuroticism.

### Author contributions

**Johanna L. Popp:** Conceptualization, Formal analysis, Funding acquisition, Methodology, Project administration, Writing - Original Draft, Visualization. **Martin Weiß:** Funding acquisition, Methodology, Writing - Review & Editing. **Joshua Faskowitz:** Data Curation, Resources, Writing - Review & Editing. **Kirsten Hilger:** Conceptualization, Funding acquisition, Supervision, Writing – Original Draft.

## Acknowledgements

Our sincere appreciation goes to the Human Connectome Project^46^, WU-Minn Consortium (Principal Investigators: David Van Essen and Kamil Ugurbil; 1U554MH091657), which was funded by the 16 NIH Institutes and Centers under the NIH Blueprint for Neuroscience Research, along with support from the McDonnell Center for Systems Neuroscience at Washington University. Further, we thank Luke Chang, Emily Finn, and Jeremy Manning for their freely available online course on naturalistic data analysis, which was an invaluable resource for this work. Lastly, we also thank David Gruskin for kindly sharing his script from a conceptually related movie rating study, which provided useful guidance.

## Additional information

The online version of this article contains supplementary material.

## Abbreviations

AnnaK(-Model) =: Anna Karenina Model
ANOVA =: analysis of variance
BOLD =: blood oxygen level dependent
DA =: dorsal attention network
DMN =: default mode network
fMRI =: functional magnetic resonance imaging
FP =: frontoparietal control network
EEG =: electroencephalography
Gauss(ian-Model) =: Gaussian Model
HCP =: Human Connectome Project
ISC =: inter-subject correlation
IS-RSA =: inter-subject representational similarity analysis
LIM =: limbic network
NN(-Model) =: Nearest Neighbors Model
SOM =: somatomotor network
TE =: echo time
TR =: repetition time
VA =: ventral attention network
VIS =: visual network

