## Supplementary for "The Neural Code of Neuroticism"

Johanna L. Popp: 0000-0003-1704-9890

Martin Weiß: 0000-0002-0569-0907

Joshua Faskowitz: 0000-0003-1814-7206

Kirsten Hilger: 0000-0003-3940-5884

* **Correspondending Authors**

Johanna L. Popp, Department of Psychology I, Würzburg University, Marcusstr. 9-11, Würzburg D-97070, Germany.

Kirsten Hilger, Department of Psychology, Vinzenz Pallotti University Vallendar, Pallottistr. 3, 56197 Vallendar, Germany.

**Abbreviations**

AnnaK(-Model) = Anna Karenina Model

ANOVA = analysis of variance

BOLD = blood oxygen level dependent

DA = dorsal attention network

DMN = default mode network

fMRI = functional magnetic resonance imaging

FP = frontoparietal control network

EEG = electroencephalography

Gauss(ian-Model) = Gaussian Model

HCP = Human Connectome Project

ISC = inter-subject correlation

IS-RSA = inter-subject representational similarity analysis

LIM = limbic network

NN(-Model) = Nearest Neighbors Model

SOM = somatomotor network

TE = echo time

TR = repetition time

VA = ventral attention network

VIS = visual network

**Supplementary Table S1**

Description of movie clips

| Movie/scan | Short name clip | Duration (min:s; frames) | Description |
|---|---|---|---|
| 1 | Two men | 04:04 (244) | In rural Australia one man, sitting down near a street, observes another man who is running past him and reflects on his potential motives. The dialogue is accented in English and subtitles are provided. |
| 1 | Bridgeville | 03:41 (221) | Individuals share their reasons for living in Bridgeville, a small town in the United States. Their reports are separated by short glimpses into community life. |
| 1 | Pockets | 03:08 (188) | Close-up portraits of individuals presenting what they carry in their pockets while explaining the personal significance of the respective items. |
| 1 | Overcome | 01:04 (64) | Inspirational montage featuring individuals who have overcome physical disabilities. |
| 1 | Testretest1 | 01:24 (84) | Sequence of short clips (1-3 seconds) showcasing a diverse range of random people, objects, and scenes. |
| 2 | Inception | 03:47 (227) | Two individuals walk around in a dream world and explore its physical and emotional characteristics. |
| 2 | Social net | 04:18 (258) | Fictional depiction of Mark Zuckerberg’s disciplinary hearing at Harvard and following events. |
| 2 | Ocean’s 11 | 04:09 (249) | Danny Ocean and his accomplices meet to plan their heist on a casino in Las Vegas. |
| 2 | Testretest2 | 01:24 (84) | Identical to testretest1. |
| 3 | Flower | 03:00 (180) | A flower is freed from its pot and gets taken around a neighborhood. |
| 3 | Hotel | 03:05 (185) | A man and a woman experience a metaphysical encounter while being in a hotel room. |
| 3 | Garden | 03:24 (204) | A woman shares information about an urban vegetable garden supporting the local community. |
| 3 | Dreary | 02:22 (142) | Compilation of dreary landscapes and abandoned buildings accompanied by spooky music. |
| 3 | Testretest3 | 01:24 (84) | Identical to testretest1. |
| 4 | Home alone | 03:52 (232) | Kevin, an eight-year-old boy, walks through the house and realizes his family has left him home alone. |
| 4 | Brokovich | 03:50 (230) | Erin Brokovich, accompanied by three young children, meets with a plaintiff and later visits a law office. |
| 4 | Star wars | 04:15 (255) | Scene at the rebel base on an icy planet where Luke gets attacked. Afterwards, Leia and Han Solo argue. |
| 4 | Testretest4 | 01:24 (84) | Identical to testretest1. |

*Note:* Movie 1 and 3 contained parts of documentaries and independent films freely available under a Creative Commons license, while Movie 2 and 4 included parts of Hollywood films[1].

The clip named “Testretest” was played at the end of each movie. More detailed information about the timing of clips and the separation into scenes for the Pilot Study is provided in Supplementary Table S2. This table was adapted from Finn & Bandettini[2]. Clip durations were determined independently and thus slightly differ.

**Supplementary Table S2**

Timing and segmentation details of movie stimuli (Main Study and Pilot Study)

| Movie | Clip | Duration (s) | From-to (min:s; frames) | Scene | From-to (min:s; frames) | Duration (s) |
|---|---|---|---|---|---|---|
| | Rest: 00:00 – 00:20 (00 – 20; 20 s) | | | | | |
| Movie 1 | Two Men | 04:04 | 00:20 – 04:24 | 1 | 00:20 – 00:53 (20 – 53) | 33 |
| 15:21 | | (244) | (20-264) | 2 | 00:53 – 01:15 (53 – 75) | 22 |
| (921) | | | | 3 | 01:15 – 01:54 (75 – 114) | 39 |
| | | | | 4 | 01:54 – 02:20 (114 – 141) | 27 |
| | | | | 5 | 02:21 – 03:06 (141 – 187) | 46 |
| | | | | 6 | 03:07 – 03:51 (187 – 232) | 45 |
| | | | | Credits | 03:52 – 04:24 (232 – 264) | 32 |
| | Rest: 04:24 – 04:44 (264 -284; 20 s) | | | | | |
| | Bridgeville | 03:41 | 04:44 – 08:25 | 7 | 04:44 – 05:14 (284 – 314) | 30 |
| | | (221) | (284 – 505) | 8 | 5:14 -5:31 (314 – 331) | 17 |
| | | | | 9 | 5:31 – 5:56 (331 – 356) | 25 |
| | | | | 10 | 5:56 – 6:52 (356 – 412) | 56 |
| | | | | 11 | 6:52 – 7:11 (412 – 431) | 19 |
| | | | | 12 | 7:11 – 7:48 (431 - 468) | 37 |
| | | | | 13 | 7:48 – 8:25 (468 – 505) | 37 |
| | Rest: 08:25 – 08:45 (505 -525; 20 s) | | | | | |
| | Pockets | 03:08 | 08:45 – 11:53 | 14 | 08:45 – 09:08 (525 – 548) | 23 |
| | | (188) | (525 – 713) | 15 | 09:08 – 09:37 (548 – 577) | 29 |
| | | | | 16 | 09:37 – 10:19 (577 – 619) | 42 |
| | | | | 17 | 10:19 – 10:56 (619 – 656) | 37 |
| | | | | 18 | 10:56 – 11:21 (656 – 681) | 25 |
| | | | | 19 | 11:21- 11:46 (681 – 706) | 25 |
| | | | | Credits | 11:46 – 11:53 (707 – 713) | 7 |
| | Rest: 11:53 – 12:13 (713 –733; 20 s) | | | | | |
| | Overcome | 01:04 (64) | 12:13 – 13:17 (733 – 797) | 20 | 12:13 – 13:17 (733 -797) | 64 |
| | Rest: 13:17 – 13:37 (797 – 817; 20 s) | | | | | |
| | Test | 01:24 | 13:37 – 15:01 | 21 | 13:37 – 14:02 (817 – 842) | 25 |
| | | (84) | (817 – 901) | 22 | 14:02 – 14:37 (842 – 877) | 35 |
| | | | | 23 | 14:37 – 15:01 (877 – 901) | 24 |
| | Rest: 15:01 – 15:21 (901 – 921; 20 s) | | | | | |
| | Rest: 00:00 – 00:20 (00 – 20; 20 s) | | | | | |
| Movie 2 | Inception | 03:47 | 00:20 – 04:07 | 1 | 00:20 -00:54 (20 – 54) | 34 |
| 15:18 | | (227) | (20 – 247) | 2 | 00:54– 1:52 (54 – 112) | 58 |
| (918) | | | | 3 | 1:52 – 2:35 (112 – 155) | 43 |
| | | | | 4 | 2:35 – 02:55 (155 – 175) | 20 |
| | | | | 5 | 02:55 – 03:42 (175 – 222) | 47 |
| | | | | 6 | 03:42 – 04:07 (222 – 247) | 25 |
| | Rest: 04:07 – 04:27 (247 -267; 20 s) | | | | | |
| | Social net | 04:18 | 04:27-08:45 | 7 | 04:27 – 05:10 (267 – 310) | 43 |
| | | (258) | (267 - 525) | 8 | 05:10 – 5:56 (310 - 356) | 46 |
| | | | | 9 | 5:56 – 06:31 (356 – 391) | 35 |
| | | | | 10 | 06:31-07:27 (391 – 447) | 56 |
| | | | | 11 | 07:27 – 07:54 (447 – 474) | 27 |
| | | | | 12 | 07:54 – 08:45 (474 -525) | 51 |
| | Rest: 08:45 – 09:05 (525 – 545; 20 s) | | | | | |
| | Ocean's 11 | 04:09 | 09:05 -13:14 | 13 | 09:05 – 09:38 (545 – 578) | 33 |
| | | (249) | (545 – 794) | 14 | 09:38 – 10:06 (578 – 606) | 28 |
| | | | | 15 | 10:06 – 10:40 (606 – 640) | 34 |
| | | | | 16 | 10:40 – 11:39 (640 – 699) | 59 |
| | | | | 17 | 11:39 – 12:29 (699 – 749) | 50 |
| | | | | 18 | 12:29 – 13:14 (749 – 794) | 45 |
| | Rest: 13:14 – 13:34 (794 – 814; 20 s) | | | | | |
| | Test | 01:24 | 13:34 – 14:58 | 19 | 13:34 – 13:59 (814 – 839) | 25 |
| | | (84) | (814 – 898) | 20 | 13:59 – 14:34 (839 –874) | 35 |
| | | | | 21 | 14:34 – 14:58 (874 – 898) | 24 |
| | Rest: 14:58 – 15:18 (898 – 918; 20 s) | | | | | |

| | | | | | | |
|---|---|---|---|---|---|---|
| | Rest: 00:00 – 00:20 (00-20; 20 s) | | | | | |
| Movie 3 | Flower | 03:00 | 00:20 – 03:20 | 1 | 00:20 – 1:04 (20 – 64) | 44 |
| 15:15 | | (180) | (20 – 200) | 2 | 1:04 – 1:47 (64 – 107) | 43 |
| (915) | | | | 3 | 1:47 – 2:31 (107 – 151) | 44 |
| | | | | 4 | 2:31 -3:20 (151 – 200) | 49 |
| | Rest 3:20 – 3:40 (200 – 220; 20 s) | | | | | |
| | Hotel | 03:05 | 03:40 – 06:45 | 5 | 3:40 – 4:33 (220 – 273) | 53 |
| | | (185) | (220 – 405) | 6 | 4:33 – 5:03 (273 – 303) | 30 |
| | | | | 7 | 5:03 – 5:33 (303 – 333) | 30 |
| | | | | 8 | 5:33 – 6:04 (333 – 364) | 31 |
| | | | | 9 | 6:04 – 6:45 (364 – 405) | 41 |
| | Rest: 6:45 – 7:05 (405 – 4:25; 20 s) | | | | | |
| | Garden | 03:24 | 7:05 – 10:29 | 10 | 7:05 – 7:31 (425 – 451) | 26 |
| | | (204) | (425 – 629) | 11 | 7:31 – 8:19 (451 – 499) | 48 |
| | | | | 12 | 8:19 – 08:57 (499 – 537) | 38 |
| | | | | 13 | 08:57 – 09:27 (537 – 567) | 30 |
| | | | | 14 | 09:27 – 09:57 (567 – 597) | 30 |
| | | | | 15 | 09:57 – 10:28 (597 – 629) | 32 |
| | Rest: 10:29 – 10:49 (629 – 649; 20 s) | | | | | |
| | Dreary | 02:22 | 10:49 – 13:11 | 16 | 10:49 – 11:23 (649 –683) | 34 |
| | | (142) | (649 – 791) | 17 | 11:23 – 12:02 (683 – 722) | 39 |
| | | | | 18 | 12:02 – 12:33 (722 – 753) | 31 |
| | | | | 19 | 12:33 – 13:11 (753 – 791) | 38 |
| | Rest: 13:11 – 13:31 (791 – 811; 20 s) | | | | | |
| | Test | 01:24 | 13:31 – 14:55 | 20 | 13:31 – 13:56 (811 – 836) | 25 |
| | | (84) | (811-895) | 21 | 13:56 – 14:31 (836 – 871) | 35 |
| | | | | 22 | 14:31 – 14:55 (871 – 895) | 24 |
| | Rest: 14:55 – 15:15 (895 – 915; 20 s) | | | | | |
| | Rest: 00:00 – 00:20 00 – 20; 20 s) | | | | | |
| Movie 4 | Home | 03:52 | 00:20 – 4:12 | 1 | 00:20 - 00:58 (20 – 58) | 38 |
| 15:01 | alone | (232) | (20 – 252) | 2 | 00:58- 01:26 (58 - 86) | 28 |
| (901) | | | | 3 | 01:26 – 02:06 (86 – 126) | 40 |
| | | | | 4 | 02:06 – 02:43 (126 – 163) | 37 |
| | | | | 5 | 02:43 – 03:14 (163 - 194) | 31 |
| | | | | 6 | 03:14 – 03:39 (194 – 219) | 25 |
| | | | | 7 | 03:39 – 4:12 (219-252) | 33 |
| | Rest: 4:12 – 4:32 (252 – 272; 20 s) | | | | | |
| | Brokovich | 03:50 | 4:32 – 08:22 | 8 | 4:32 – 05:06 (272 – 306) | 34 |
| | | (230) | (272 – 502) | 9 | 05:06-05:44 (306 – 344) | 38 |
| | | | | 10 | 05:44 -06:26 (344 – 386) | 42 |
| | | | | 11 | 06:26 – 07:03 (386 – 423) | 37 |
| | | | | 12 | 07:03- 07:41 (423 – 461) | 38 |
| | | | | 13 | 07:41 – 08:22 (461 – 502) | 41 |
| | Rest: 08:22 – 08:42 (502 -522; 20 s) | | | | | |
| | Star wars | 04:15 | 08:42 – 12:57 | 14 | 08:42 – 09:13 (522 – 553) | 31 |
| | | (255) | (522 – 777) | 15 | 09:13 – 09:41 (553 – 581) | 28 |
| | | | | 16 | 09:41 - 10:27 (581 – 627) | 46 |
| | | | | 17 | 10:27 – 11:09 (627 – 669) | 42 |
| | | | | 18 | 11:09 – 12:12 (669 – 732) | 63 |
| | | | | 19 | 12:12 – 12:57 (732 -777) | 45 |
| | Rest: 12:57 – 13:17 (777 - 797; 20 s) | | | | | |
| | Test | 01:24 | 13:17 – 14:41 | 20 | 13:17 – 13:42 (797 - 822) | 25 |
| | | (84) | (797 – 881) | 21 | 13:42 -14:17 (822 – 857) | 35 |
| | | | | 22 | 14:17 – 14:41 (857 – 881) | 24 |
| | Rest: 14:41 – 15:01 (881 – 901; 20 s) | | | | | |

*Note:* Clips contained in the four movies shown to participants of the Main Study ($N$ = 174) were further segmented into scenes of approximately 30 seconds and rated for their trait-relevance by participants in the Pilot Study ($N$ = 86). Scene durations varied to preserve the narrative flow of the movie content and minor adjustments (e.g., trimming frames) were made to ensure smooth scene transitions. Group 1 ($N$ = 41) rated scenes from Movies 1 and 2, while Group 2 ($N$ = 45) rated scenes from Movies 3 and 4. Scenes identified as trait-relevant are marked in red while scenes identified as trait-irrelevant are marked in green. Rest periods and credits were excluded in the Pilot Study.

**Supplementary Table S3**

Statements tailored to assess the trait-relevance of scenes presented in the Pilot Study

| **Neuroticism facet** | **Statement** |
|---|---|
| Regular-coded | |
| Anxiety | This scene made me feel tense. |
| Angry hostility | This scene made me feel frustrated. |
| Self-consciousness | This scene made me feel dissatisfied with myself. |
| Reverse-coded | |
| Depression | This scene made me feel happy. |
| Impulsiveness | This scene made me feel safe and secure |
| Vulnerability | This scene made me feel confident in myself and my abilities. |

*Note:* After each scene with a duration of approximately 30 seconds, participants were asked for their level of agreement with each of the above presented statements, each specifically addressing one facet of neuroticism, on a five-point Likert scale ranging from 0 (strongly disagree) to 4 (strongly agree). Three of the six statements were reverse-coded to assess the expression of negative neuroticism-related emotions and their positive counterparts, aiming to avoid response bias and foster a comfortable atmosphere.

**Supplementary Table S4**

Brain-trait representational similarity at the whole-brain and network-level during trait-relevant scenes in the main sample (*N* = 89) and the replication sample (*N* = 85) operationalized with all models of trait similarity

| Trait-relevant movie scenes | | | | | | | | | | | | |
|---|---|---|---|---|---|---|---|---|---|---|---|---|
| | **Main sample (*N* = 89)** | | | | | | **Replication sample (*N* = 85)** | | | | | |
| **Network (YEO 7)** | **NN** | | **AnnaK** | | **Gauss** | | **NN** | | **AnnaK** | | **Gauss** | |
| | ***t*** | ***p*** | ***t*** | ***p*** | ***t*** | ***p*** | ***t*** | ***p*** | ***t*** | ***p*** | ***t*** | ***p*** |
| All (200 regions) | .94 | .351 | -20.35 | <.001 | 2.79 | .006 | -2.06 | .040 | -18.15 | <.001 | -2.32 | .022 |
| DMN (50 regions) | .67 | .506 | -10.04 | <.001 | -.75 | .454 | -.39 | .700 | -11.70 | <.001 | -.15 | .881 |
| FP (25 regions) | .26 | .793 | -5.10 | <.001 | .33 | .747 | -.44 | .662 | -8.87 | <.001 | -.07 | .941 |
| LIM (10 regions) | 1.53 | .161 | -3.43 | .008 | 1.22 | .255 | -1.44 | .185 | -2.58 | .030 | -.88 | .402 |
| VA (22 regions) | .13 | .894 | -8.30 | <.001 | 1.51 | .146 | -.43 | .670 | -5.68 | <.001 | -.86 | .399 |
| DA (31 regions) | .56 | .581 | -11.08 | <.001 | 1.24 | .225 | -1.16 | .257 | -6.86 | <.001 | -1.30 | .205 |
| SOM (37 regions) | -.32 | .748 | -7.28 | <.001 | 2.48 | .018 | -.29 | .771 | -4.14 | <.001 | -1.01 | .318 |
| VIS (25 regions) | .14 | .894 | -15.68 | <.001 | .55 | .585 | -2.80 | .010 | -16.58 | <.001 | -2.55 | .017 |

*Note:* Model-specific IS-RSA correlation coefficients computed with brain region-specific activity during trait-relevant movie scenes in all 200 brain regions and subsets of these brain regions belonging to each of the seven Yeo functional networks[3] were tested for significant brain-trait representational similarity by performing one-sample *t*-tests. NN = NN-Model; AnnaK = AnnaK-Model; Gauss = Gaussian-Model; DMN = default mode network; FP = frontoparietal control network; LIM = limbic network; VA = ventral attention/salience network; DA = dorsal attention network; SOM = somatomotor network; VIS = visual network.

**Supplementary Table S5**

Brain-trait representational similarity at the whole-brain and network-level during trait-irrelevant movie scenes in the main sample ($N$ = 89) and the replication sample ($N$ = 85) operationalized with all models of trait similarity

| Trait-irrelevant movie scenes | | | | | | | | | | | | |
|---|---|---|---|---|---|---|---|---|---|---|---|---|
| | **Main sample (*N* = 89)** | | | | | | **Replication sample (*N* = 85)** | | | | | |
| **Network (YEO 7)** | **NN** | | **AnnaK** | | **Gauss** | | **NN** | | **AnnaK** | | **Gauss** | |
| | ***t*** | ***p*** | ***t*** | ***p*** | ***t*** | ***p*** | ***t*** | ***p*** | ***t*** | ***p*** | ***t*** | ***p*** |
| All (200 regions) | -6.62 | <.001 | -9.10 | <.001 | -7.03 | <.001 | 7.39 | <.001 | -10.70 | <.001 | 5.71 | <.001 |
| DMN (50 regions) | -6.73 | <.001 | -3.74 | <.001 | -7.82 | <.001 | 8.92 | <.001 | -8.99 | <.001 | 7.40 | <.001 |
| FP (25 regions) | -3.42 | .002 | -4.22 | <.001 | -3.89 | .001 | 5.53 | <.001 | -5.60 | <.001 | 5.54 | <.001 |
| LIM (10 regions) | -.41 | .692 | -.80 | .444 | -.56 | .586 | 2.52 | .033 | -.90 | .390 | 2.95 | .016 |
| VA (22 regions) | -2.10 | .048 | -.87 | .397 | -1.96 | .064 | 1.23 | .233 | -3.27 | .004 | .57 | .577 |
| DA (31 regions) | -2.15 | .040 | -3.21 | .003 | -1.94 | .062 | 2.47 | .019 | -7.68 | <.001 | 2.45 | .020 |
| SOM (37 regions) | -.58 | .569 | -3.13 | .003 | -.39 | .698 | 2.42 | .021 | 1.84 | .074 | .96 | .345 |
| VIS (25 regions) | -2.46 | .021 | -17.99 | <.001 | -2.61 | .015 | -2.31 | .030 | -8.31 | <.001 | -2.51 | .019 |

*Note:* Model-specific IS-RSA correlation coefficients computed with brain region-specific activity during trait-irrelevant movie scenes in all 200 brain regions and subsets of these brain regions belonging to each of the seven Yeo functional networks[3] were tested for significant brain-trait representational similarity by performing one-sample $t$-tests. NN = NN-Model; AnnaK = AnnaK-Model; Gauss = Gaussian-Model; DMN = default mode network; FP = frontoparietal control network; LIM = limbic network; VA = ventral attention/salience network; DA = dorsal attention network; SOM = somatomotor network; VIS = visual network.

**Supplementary Table S6**

Brain-trait representational similarity at the brain region-specific level during trait-relevant movie scenes in the main sample (*N* = 89) and the replication sample (*N* = 85) operationalized with all models of trait similarity

| **Trait-relevant movie scenes** | | | | | | | |
|---|---|---|---|---|---|---|---|
| **Main sample (*N* = 89)** | | | | **Replication sample (*N* = 85)** | | | |
| **NN-Model** | | | | | | | |
| | **Name** | ***r*** | ***p*** | | **Name** | ***r*** | ***p*** |
| 1 | LH_DefaultC_RSC | -.06 | .035 | 1 | LH_DefaultA_pCun | -.06 | .044 |
| 2 | LH_LimbicB_OFC_1 | .04 | .042 | 2 | LH_SomMotB_3 | -.05 | .012 |
| 3 | LH_SalVentAttnA_Ins_2 | -.05 | .017 | 3 | RH_DorsAttnB_SPL_1 | -.07 | .039 |
| | | | | 4 | RH_SomMotB_4 | -.04 | .048 |
| | | | | 5 | RH_SomMotB_ST_1 | .08 | .018 |
| **AnnaK-Model** | | | | | | | |
| | **Name** | ***r*** | ***p*** | | **Name** | ***r*** | ***p*** |
| 1 | LH_TempPar_Temp_3 | -.23 | .015 | 1 | LH_DefaultB_IPL_2 | -.20 | .017 |
| 2 | LH_DefaultA_PCC | -.16 | .018 | 2 | LH_DefaultB_PFCd_2 | -.16 | .035 |
| 3 | LH_DefaultA_PFCm_3 | -.11 | .038 | 3 | LH_DefaultB_PFCv_2 | -.19 | .020 |
| 4 | LH_ContC_PCC | -.15 | .009 | 4 | LH_DefaultB_TempPole | -.20 | .028 |
| 5 | LH_SalVentAttnA_FrMed_2 | -.08 | .030 | 5 | LH_DefaultB_Temp_1 | -.19 | .029 |
| 6 | LH_SomMotB_Ins_2 | -.07 | .042 | 6 | LH_DefaultB_Temp_2 | -.21 | .028 |
| 7 | LH_VisCent_ExStr_5 | -.18 | .042 | 7 | LH_DefaultA_pCun | -.15 | .039 |
| **8** | **RH_SalVentAttnB_PFCv_2** | **-.15** | **.026** | 8 | LH_ContA_IPS_2 | -.13 | .048 |
| 9 | RH_ContB_IPL_2 | -.13 | .008 | 9 | LH_ContA_PFCl_1 | -.19 | .011 |
| 10 | RH_DefaultA_PFCm_1 | -.12 | .037 | 10 | LH_LimbicB_OFC_2 | -.13 | .025 |
| 11 | RH_ContA_PFCl_1 | -.19 | .036 | 11 | LH_DorsAttnA_SPL_2 | -.18 | .047 |
| 12 | RH_LimbicA_Temp | -.04 | .046 | 12 | LH_DorsAttnA_TempOcc_4 | -.20 | .026 |
| 13 | RH_SalVentAttnB_FrMed_1 | -.13 | .050 | 13 | LH_SomMotA_3 | -.12 | .006 |
| 14 | RH_SalVentAttnB_FrMed_2 | -.12 | .049 | 14 | RH_DefaultB_PFCd_1 | -.14 | .024 |
| 15 | RH_SalVentAttnA_IPL | -.13 | .002 | 15 | RH_DefaultB_PFCd_2 | -.19 | .010 |
| 16 | RH_DorsAttnA_TempOcc_1 | -.18 | .044 | 16 | RH_DefaultB_PFCv_1 | -.20 | .007 |
| 17 | RH_DorsAttnA_TempOcc_3 | -.22 | .027 | **17** | **RH_SalVentAttnB_PFCv_2** | **-.18** | **.006** |
| 18 | RH_SomMotB_Ins_1 | -.09 | .011 | 18 | RH_DefaultB_Temp_1 | -.09 | .034 |
| 19 | RH_SomMotB_Ins_3 | -.08 | .018 | 19 | RH_DefaultA_pCun | -.11 | .036 |
| 20 | RH_VisCent_ExStr_2 | -.20 | .037 | 20 | RH_ContC_PCC | -.12 | .037 |
| | | | | 21 | RH_ContC_pCun_2 | -.13 | .024 |
| | | | | 22 | RH_ContB_FPole | -.15 | .017 |
| | | | | 23 | RH_ContB_PFCv | -.17 | .024 |
| | | | | 24 | RH_LimbicA_TempPole | -.14 | .017 |
| | | | | 25 | RH_DorsAttnA_SPL_1 | .17 | .048 |
| | | | | 26 | RH_DorsAttnA_SPL_2 | -.20 | .019 |
| | | | | 27 | RH_SomMotA_3 | -.13 | .004 |
| | | | | 28 | RH_SomMotA_4 | -.11 | .040 |
| **Gaussian-Model** | | | | | | | |
| | **Name** | ***r*** | ***p*** | | **Name** | ***r*** | ***P*** |
| | | | | 1 | LH_SomMotB_3 | -.06 | .032 |
| | *N/A* | | | 2 | RH_SomMotB_Ins_1 | -.07 | .033 |
| | | | | 3 | RH_SomMotB_ST_1 | .14 | .024 |

*Note:* Brain regions with significant brain-trait representational similarity, as identified via non-parametric permutation testing (Mantel-test; uncorrected threshold of $p < 0.05$), during trait-relevant movie scenes. Please note that no brain region remained significant after false discovery rate (FDR) correction for multiple comparisons. Brain regions showing significant brain-trait representational similarity in the main and the replication sample are highlighted in bold.

**Supplementary Table S7**

Brain-trait representational similarity at the brain region-specific level during trait-irrelevant movie scenes in the main sample (*N* = 89) and the replication sample (*N* = 85) operationalized with all models of trait similarity

| **Trait-irrelevant movie scenes** | | | | | | | |
|---|---|---|---|---|---|---|---|
| **Main sample (*N* = 89)** | | | | **Replication sample (*N* = 85)** | | | |
| **NN-Model** | | | | | | | |
| | **Name** | ***r*** | ***p*** | | **Name** | ***r*** | ***p*** |
| 1 | LH_DorsAttnB_SPL_2 | -.07 | .024 | 1 | LH_DefaultA_PFCd_2 | .04 | .047 |
| 2 | LH_SomMotB_Ins_3 | -.06 | .048 | 2 | LH_SomMotA_7 | .05 | .042 |
| 3 | RH_DorsAttnA_SPL_1 | -.08 | .012 | 3 | LH_SomMotA_9 | .06 | .005 |
| | | | | 4 | RH_LimbicB_OFC_1 | .04 | .048 |
| **AnnaK-Model** | | | | | | | |
| | **Name** | ***r*** | ***p*** | | **Name** | ***r*** | ***p*** |
| 1 | RH_TempPar_Temp_2 | -.19 | .038 | 1 | LH_DefaultC_IPL | -.17 | .034 |
| 2 | RH_DefaultA_PFCd_1 | -.08 | .023 | 2 | LH_DefaultC_RSC | -.18 | .037 |
| 3 | RH_SomMotA_4 | -.08 | .034 | 3 | LH_DefaultA_IPL_1 | -.12 | .029 |
| | | | | 4 | LH_DefaultA_PCC | -.10 | .049 |
| | | | | 5 | LH_DefaultA_pCun | -.13 | .020 |
| | | | | 6 | LH_SalVentAttnA_FrMed_1 | -.12 | .039 |
| | | | | 7 | LH_DorsAttnA_SPL_2 | -.20 | .023 |
| | | | | 8 | LH_DorsAttnA_TempOcc_4 | -.24 | .010 |
| | | | | 9 | LH_SomMotA_5 | .08 | .024 |
| | | | | 10 | RH_SalVentAttnB_IPL | -.20 | .028 |
| | | | | 11 | RH_DefaultB_PFCd_2 | -.14 | .028 |
| | | | | 12 | RH_DefaultB_PFCv_1 | -.137 | .041 |
| | | | | 13 | RH_SalVentAttnB_PFCv_2 | -.12 | .031 |
| | | | | 14 | RH_DefaultB_TempPole | -.12 | .012 |
| | | | | 15 | RH_DefaultB_Temp_1 | -.09 | .013 |
| | | | | 16 | RH_DefaultA_PFCd_3 | -.18 | .002 |
| | | | | 17 | RH_DefaultA_PFCm_2 | -.13 | .047 |
| | | | | 18 | RH_ContB_IPL_1 | -.17 | .016 |
| | | | | 19 | RH_ContB_PFCv | -.14 | .031 |
| | | | | 20 | RH_ContA_PFCl_1 | -.19 | .046 |
| | | | | 21 | RH_DorsAttnA_TempOcc_2 | -.19 | .031 |
| | | | | 22 | RH_SomMotA_10 | .08 | .009 |
| | | | | 23 | RH_SomMotA_5 | .09 | .013 |
| | | | | 24 | RH_SomMotA_6 | .12 | .001 |
| | | | | 25 | RH_SomMotA_7 | .12 | .010 |
| **Gaussian-Model** | | | | | | | |
| | **Name** | ***r*** | ***p*** | | **Name** | ***r*** | ***p*** |
| 1 | LH_DorsAttnB_SPL_2 | -.11 | .038 | 1 | LH_DefaultA_PFCd_2 | .07 | .018 |
| 2 | RH_ContB_Temp | -.10 | .042 | 2 | LH_SomMotA_9 | .06 | .040 |
| 3 | RH_DorsAttnA_SPL_1 | -.13 | .023 | 3 | RH_ContC_pCun_2 | .06 | .042 |

*Note:* Brain regions with significant brain-trait representational similarity, as identified via non-parametric permutation testing (Mantel-test; uncorrected threshold $p < 0.05$), during trait-irrelevant movie scenes. Please note that no brain region remained significant after FDR correction for multiple comparisons. No brain region showed significant brain-trait representational similarity in the main sample and the replication sample.

**Supplementary Table S8**

Brain-trait representational similarity at the whole-brain and network-level during all movie scenes in the main sample ($N$ = 89) and the replication sample ($N$ = 85) operationalized with all models of trait similarity

| All movie scenes | | | | | | | | | | | | |
|---|---|---|---|---|---|---|---|---|---|---|---|---|
| | **Main sample (*N* = 89)** | | | | | | **Replication sample (*N* = 85)** | | | | | |
| **Network (YEO 7)** | **NN** | | **AnnaK** | | **Gauss** | | **NN** | | **AnnaK** | | **Gauss** | |
| | ***t*** | ***p*** | ***t*** | ***p*** | ***t*** | ***p*** | ***t*** | ***p*** | ***t*** | ***p*** | ***t*** | ***p*** |
| All (200 regions) | -.44 | .660 | -21.19 | <.001 | .28 | .781 | -.77 | .441 | -17.99 | <.001 | -1.95 | .052 |
| DMN (50 regions) | -.41 | .683 | -10.65 | <.001 | -.54 | .595 | .75 | .460 | -15.15 | <.001 | .31 | .759 |
| FP (25 regions) | .14 | .891 | -8.58 | <.001 | -.63 | .534 | -.45 | .656 | -10.47 | <.001 | -.50 | .621 |
| LIM (10 regions) | 1.07 | .312 | -3.30 | .009 | .73 | .483 | -.75 | .473 | -2.55 | .031 | -.47 | .651 |
| VA (22 regions) | .60 | .554 | -5.52 | <.001 | .87 | .396 | .13 | .896 | -5.34 | <.001 | -.56 | .583 |
| DA (31 regions) | -.86 | .399 | -8.24 | <.001 | -.41 | .688 | -.70 | .492 | 9.34 | <.001 | -1.19 | .245 |
| SOM (37 regions) | -.47 | .643 | -7.47 | <.001 | 1.07 | .291 | .82 | .419 | -1.75 | .088 | -.22 | .827 |
| VIS (25 regions) | -.47 | .641 | -25.09 | <.001 | -.08 | .935 | -3.11 | .005 | -13.16 | <.001 | -3.46 | <.001 |

*Note:* Model-specific IS-RSA correlation coefficients computed with brain region-specific activity during all movie scenes in all 200 brain regions and subsets of these brain regions belonging to each of the seven Yeo networks[3] were tested for significant brain-trait representational similarity by performing one-sample *t*-tests. NN = NN-Model; AnnaK = AnnaK-Model; Gauss = Gaussian-Model; DMN = default mode network; FP = frontoparietal control network; LIM = limbic network; VA = ventral attention/salience network; DA = dorsal attention network; SOM = somatomotor network; VIS = visual network.

**Supplementary Table S9**

Brain-trait representational similarity at the brain region-specific level during all movie scenes in the main sample ($N$ = 89) and replication sample ($N$ = 85) operationalized with all models of trait similarity

| All movie scenes | | | | | | | |
|---|---|---|---|---|---|---|---|
| **Main sample (*N* = 89)** | | | | **Replication sample (*N* = 85)** | | | |
| **NN-Model** | | | | | | | |
| | **Name** | ***r*** | ***p*** | | **Name** | ***r*** | ***p*** |
| | *N/A* | | | 1 | RH_ContB_PFCl | .08 | .030 |
| **AnnaK-Model** | | | | | | | |
| | **Name** | ***r*** | ***p*** | | **Name** | ***r*** | ***p*** |
| 1 | LH_TempPar_Temp_3 | -.24 | .016 | 1 | LH_DefaultB_IPL_1 | -.16 | .048 |
| 2 | LH_SomMotA_10 | -.07 | .034 | 2 | LH_DefaultB_IPL_2 | -.21 | .021 |
| 3 | LH_SomMotA_6 | -.12 | .012 | 3 | LH_DefaultB_PFCv_3 | -.15 | .025 |
| 4 | LH_SomMotA_7 | -.12 | .031 | 4 | LH_DefaultB_Temp_1 | -.18 | .046 |
| 5 | RH_TempPar_Temp_1 | -.19 | .039 | 5 | LH_DefaultA_IPL_1 | -.18 | .029 |
| 6 | RH_TempPar_Temp_2 | -.20 | .025 | 6 | LH_DefaultA_PCC | -.22 | .005 |
| 7 | RH_TempPar_Temp_3 | -.19 | .049 | 7 | LH_DefaultA_PFCm_1 | -.17 | .021 |
| 8 | RH_DefaultA_PFCd_1 | -.12 | .012 | 8 | LH_DefaultA_PFCm_3 | -.14 | .026 |
| 9 | RH_SalVentAttnB_FrMed_1 | -.15 | .029 | 9 | LH_ContA_IPS_1 | -.17 | .033 |
| 10 | RH_DorsAttnB_SPL_2 | -.22 | .017 | 10 | LH_ContA_PFCl_1 | -.19 | .022 |
| 11 | **RH_DorsAttnA_TempOcc_3** | -.22 | .021 | 11 | LH_DorsAttnA_IPS | -.19 | .037 |
| | | | | 12 | LH_DorsAttnA_SPL_2 | -.22 | .026 |
| | | | | 13 | LH_DorsAttnA_TempOcc_4 | -.28 | .006 |
| | | | | 14 | LH_SomMotA_2 | -.11 | .048 |
| | | | | 15 | LH_SomMotA_4 | -.13 | .050 |
| | | | | 16 | RH_DefaultB_PFCd_1 | -.17 | .031 |
| | | | | 17 | RH_DefaultB_PFCd_2 | -.23 | .009 |
| | | | | 18 | RH_DefaultB_PFCv_1 | -.26 | .003 |
| | | | | 19 | RH_SalVentAttnB_PFCv_1 | -.20 | .012 |
| | | | | 20 | RH_SalVentAttnB_PFCv_2 | -.25 | .002 |
| | | | | 21 | RH_DefaultB_Temp_1 | -.15 | .010 |
| | | | | 22 | RH_DefaultA_PCC | -.15 | .013 |
| | | | | 23 | RH_DefaultA_PFCd_3 | -.17 | .048 |
| | | | | 24 | RH_DefaultA_PFCm_2 | -.18 | .041 |
| | | | | 25 | RH_DefaultA_pCun | -.16 | .021 |
| | | | | 26 | RH_ContC_PCC | -.23 | .004 |
| | | | | 27 | RH_ContC_pCun_1 | -.15 | .022 |
| | | | | 28 | RH_ContC_pCun_2 | -.16 | .018 |
| | | | | 29 | RH_ContB_FPole | -.21 | .008 |
| | | | | 30 | RH_ContB_IPL_1 | -.20 | .020 |
| | | | | 31 | RH_ContB_PFCv | -.21 | .011 |
| | | | | 32 | RH_ContA_PFCl_1 | -.22 | .027 |
| | | | | 33 | RH_DorsAttnA_SPL_2 | -.22 | .020 |
| | | | | 34 | RH_DorsAttnA_TempOcc_2 | -.21 | .031 |
| | | | | 35 | **RH_DorsAttnA_TempOcc_3** | -.21 | .038 |
| | | | | 36 | RH_SomMotA_2 | -.12 | .039 |
| | | | | 37 | RH_SomMotA_4 | -.13 | .050 |
| **Gaussian-Model** | | | | | | | |
| | **Name** | ***r*** | ***p*** | | **Name** | ***r*** | ***p*** |
| 1 | RH_ContA_PrCv_1 | .105 | .0374 | 1 | RH_ContB_PFCl | .133 | .040 |

*Note:* Brain regions with significant brain-trait representational similarity, as identified via non-parametric permutation testing (Mantel-test; uncorrected threshold $p < 0.05$), during all movie scenes. Please note that no brain region remained significant after FDR correction for multiple comparisons. Brain regions who showed significant brain-trait representational similarity in the main and the replication sample are highlighted in bold.

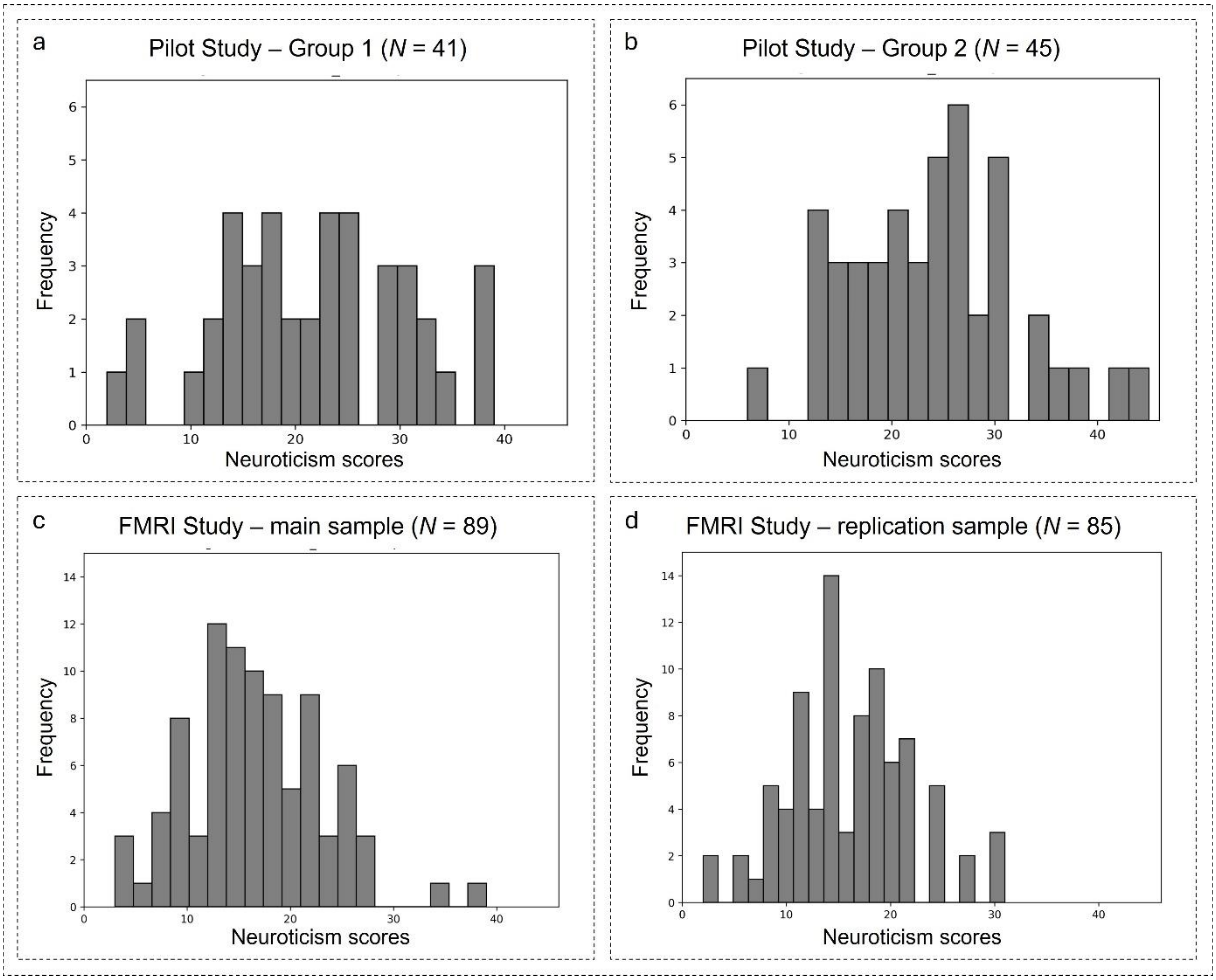


**Supplementary Fig. S1. Distribution of neuroticism scores from the NEO-FFI in the Pilot Study and the Main Study. (a)** In Group 1 of the Pilot Study ($N$ = 41), neuroticism scores ranged from 2 - 39 ($M$ = 21.7; $SD$ = 9.2) and a Shapiro Wilk test indicated that the distribution did not significantly deviate from normality ($W(41)$ = 0.978, $p$ = 0.591). **(b)** In Group 2 of the Pilot Study ($N$ = 45), neuroticism scores ranged from 6 - 45 ($M$ = 23.9; $SD$ = 8.4) and the distribution did not significantly deviate from normality ($W(45)$ = 0.985, $p$ = 0.811). **(c)** In the main sample of the Main Study ($N_{main}$ = 89), neuroticism scores ranged from 3 - 39 ($M$ = 16.5; $SD$ = 6.6) and the distribution did not significantly deviate from normality ($W(89)$ = 0.980, $p$ = 0.177). **(d)** In the replication sample of the Main Study ($N_{replication}$ = 85), neuroticism scores ranged from 2 - 31 ($M$ = 16.2; $SD$ = 6.0) and the distribution did not significantly deviate from normality ($W(85)$ = 0.987, $p$ = 0.565).

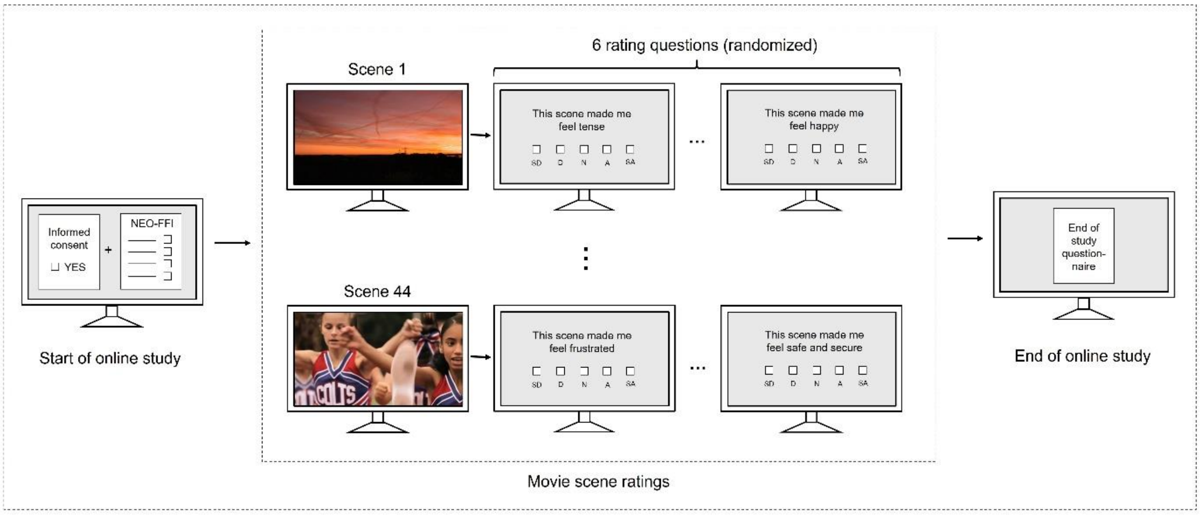


**Supplementary Fig. S2. Pilot study paradigm.** To assess the trait-relevance of the movie stimuli presented to participants of the Main Study (*N* = 174), clips contained in the four movies were further segmented into ~30-second scenes. These scenes were rated for their potential to evoke neuroticism-related emotions by participants in an independent Pilot study (*N* = 86), conducted online. To keep session length below one hour, participants were assigned to two conditions: Group 1 (*N* = 41) rated 44 scenes from Movies 1 and 2, while Group 2 (*N* = 45) rated 44 scenes from Movies 3 and 4. The study was structured as follows: After being introduced to purpose and plan of the study, participants provided informed consent, demographic information (age, gender, highest level of education, years of education, and occupation), and completed an online version of the NEO-FFI questionnaire. Then, the scenes were played in consecutive order, each followed by six statements covering the six facets of neuroticism (anxiety, angry hostility, self-consciousness, depression, impulsiveness, and vulnerability). Participants provided their level of agreement with each statement on a five-point Likert scale ranging from 0 (strongly disagree) to 4 (strongly agree). Lastly, an end-of-study questionnaire - also asking for self-assessed level of attention - was completed.

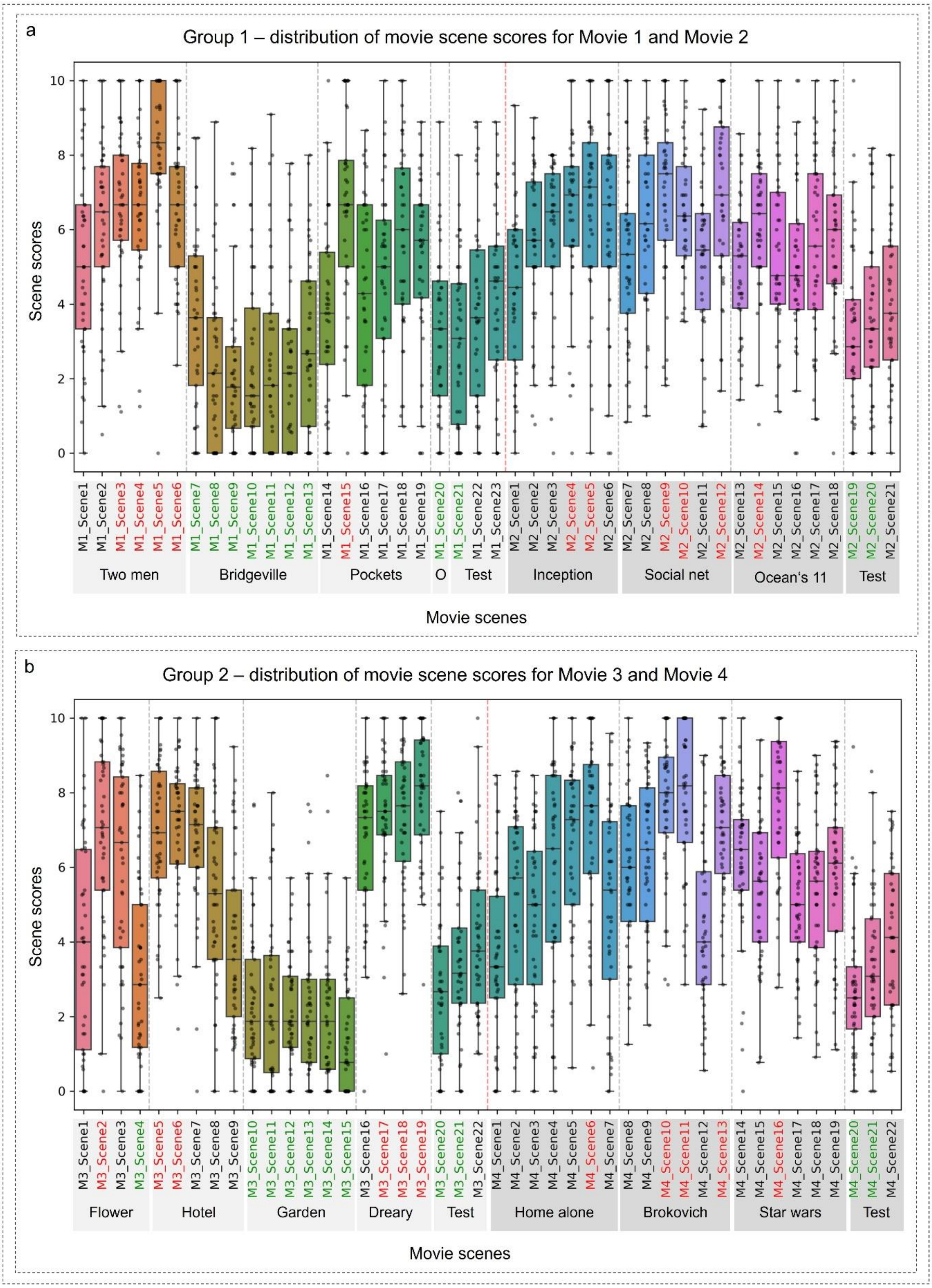
a
Group 1 – distribution of movie scene scores for Movie 1 and Movie 2
Scene scores
0
2
4
6
8
10
M1_Scene1
M1_Scene2
M1_Scene3
M1_Scene4
M1_Scene5
M1_Scene6
M1_Scene7
M1_Scene8
M1_Scene9
M1_Scene10
M1_Scene11
M1_Scene12
M1_Scene13
M1_Scene14
M1_Scene15
M1_Scene16
M1_Scene17
M1_Scene18
M1_Scene19
M1_Scene20
M1_Scene21
M1_Scene22
M1_Scene23
M2_Scene1
M2_Scene2
M2_Scene3
M2_Scene4
M2_Scene5
M2_Scene6
M2_Scene7
M2_Scene8
M2_Scene9
M2_Scene10
M2_Scene11
M2_Scene12
M2_Scene13
M2_Scene14
M2_Scene15
M2_Scene16
M2_Scene17
M2_Scene18
M2_Scene19
M2_Scene20
M2_Scene21
Two men
Bridgeville
Pockets
O
Test
Inception
Social net
Ocean's 11
Test
Movie scenes
b
Group 2 – distribution of movie scene scores for Movie 3 and Movie 4
Scene scores
0
2
4
6
8
10
M3_Scene1
M3_Scene2
M3_Scene3
M3_Scene4
M3_Scene5
M3_Scene6
M3_Scene7
M3_Scene8
M3_Scene9
M3_Scene10
M3_Scene11
M3_Scene12
M3_Scene13
M3_Scene14
M3_Scene15
M3_Scene16
M3_Scene17
M3_Scene18
M3_Scene19
M3_Scene20
M3_Scene21
M3_Scene22
M4_Scene1
M4_Scene2
M4_Scene3
M4_Scene4
M4_Scene5
M4_Scene6
M4_Scene7
M4_Scene8
M4_Scene9
M4_Scene10
M4_Scene11
M4_Scene12
M4_Scene13
M4_Scene14
M4_Scene15
M4_Scene16
M4_Scene17
M4_Scene18
M4_Scene19
M4_Scene20
M4_Scene21
M4_Scene22
Flower
Hotel
Garden
Dreary
Test
Home alone
Brokovich
Star wars
Test
Movie scenes

**Supplementary Fig. S3. Movie scene scores. (a)** Distribution of scene scores in Group 1 (*N* = 41), where participants rated a total of 44 scenes from Movies 1 and 2 (Supplementary Tables S1-2) based on their potential to evoke emotions related to neuroticism. After each scene, participants responded to six statements reflecting the six facets of neuroticism (anxiety, angry hostility, self-consciousness, depression, impulsiveness, and vulnerability), using a five-point Likert scale. For each participant, a single score (the scene score) per scene was computed by summing their provided responses, while reverse-coding relevant items. The 44 participant-specific scene scores were scaled to a range from 1 - 10 within each participant to account for individual differences in response tendencies. To obtain an overall score for each scene, scaled scores were averaged across all participants. Scenes with average scene scores in the top quartile (11 scenes, marked in red) were classified as trait-relevant, while those in the bottom quartile (11 scenes, marked in green) were classified as trait-irrelevant. **(b)** Distribution of scene scores in Group 2 (*N* = 45), where participants rated a total of 44 movie scenes from Movies 3 and 4. Scenes were rated and scored using the same procedure as in Group 1. Boxplots show the median (line), interquartile range (box spanning from 25$^{th}$ to 75$^{th}$ percentile), and whiskers extend to datapoints within 1.5 x inter quartile range.

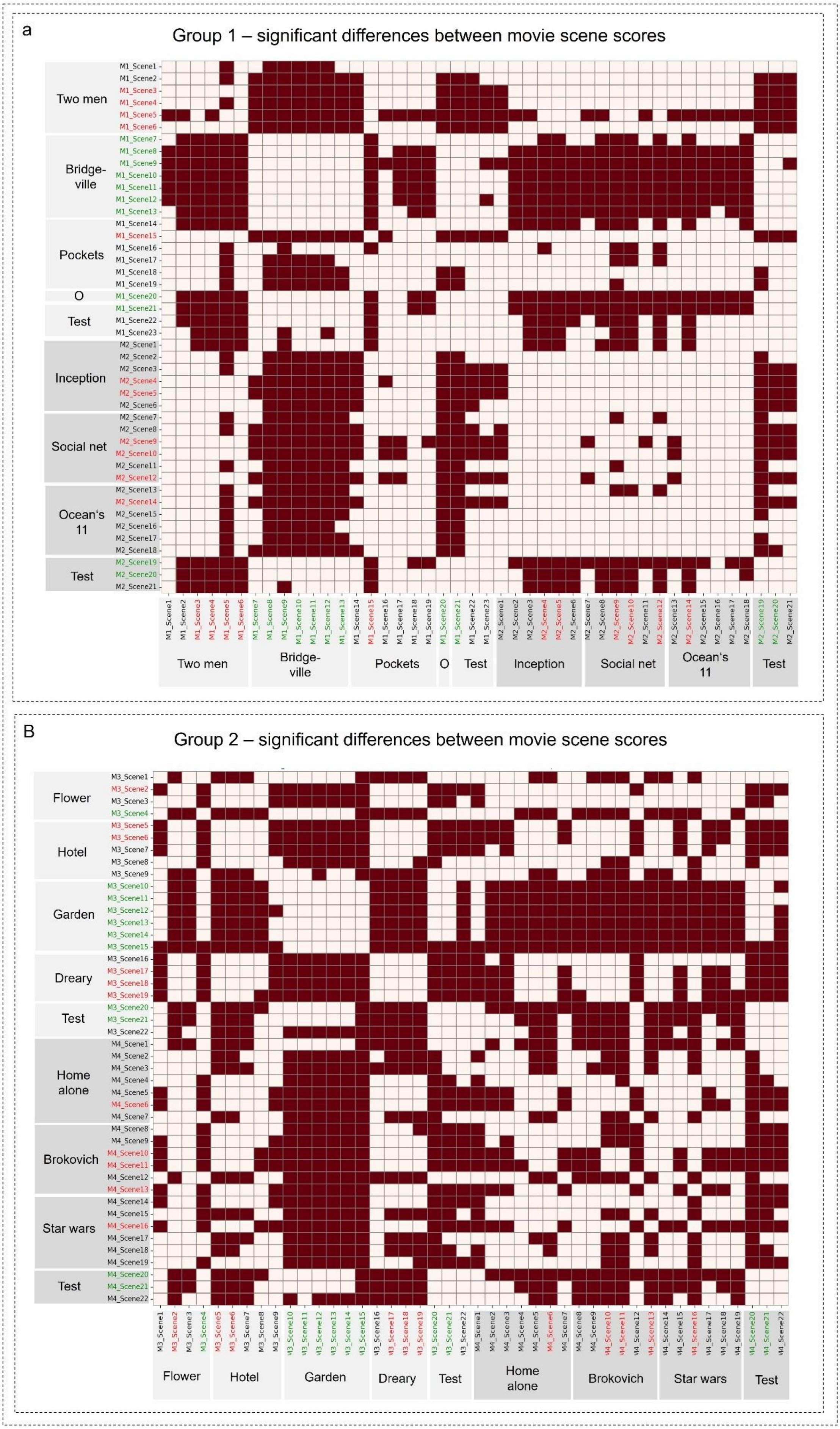
a
Group 1 – significant differences between movie scene scores
Two men
Bridge-ville
Pockets
O
Test
Inception
Social net
Ocean's 11
Test
B
Group 2 – significant differences between movie scene scores
Flower
Hotel
Garden
Dreary
Test
Home alone
Brokovich
Star wars
Test

**Supplementary Fig. S4. Significant differences between scene scores in the online study**. **(a)** A one-way repeated-measures ANOVA revealed a significant main effect of scene on participant ratings (i.e., scene scores) in Movies 1 and 2 rated by Group 1: $F(43, 1720) = 23.51$, $p < 0.001$, partial $\eta^2 = 0.31$. Post-hoc pairwise comparisons showcased multiple significant differences between individual scenes ($p < 0.05$, Holm-adjusted), which are highlighted in red. This pattern confirms that all trait-relevant scenes significantly differed from all trait-irrelevant scenes in their participant ratings (i.e., they were higher). **(b)** A one-way repeated-measures ANOVA also revealed a significant main effect of scene on participant ratings in Movies 3 and 4 rated by Group 2: $F(43, 1892) = 39.35$, $p < 0.001$, partial $\eta^2 = 0.44$. Post-hoc pairwise comparisons revealed multiple significant differences between individual scenes ($p < 0.05$, Holm-adjusted), which are highlighted in red. Again, this pattern confirmed that all trait-relevant scenes significantly differed from all trait-irrelevant scenes in their participant ratings (i.e., they were higher).

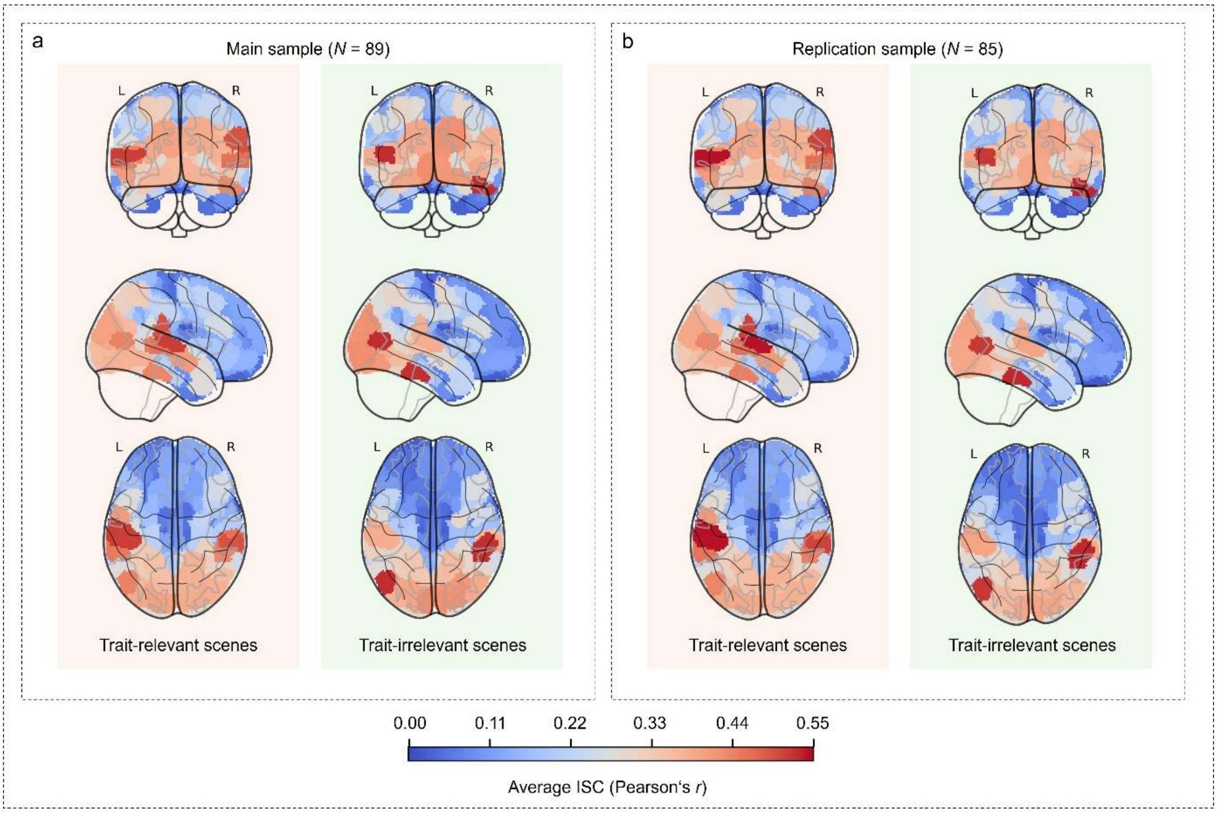


**Supplementary Fig. S5. Group-general, whole-brain pattern of similarity in brain activity (neural synchrony) during trait-relevant vs. trait-irrelevant scenes. (a)** Average neural synchrony in the main sample (*N* = 89) was computed by first correlating brain region-specific activity time courses (Pearson correlation; ISC) from respective movie scenes (either trait-relevant or trait-irrelevant) for each pair of participants, while subsequently averaging region-specific ISC correlation coefficients across all participant pairs. **(b)** Average neural synchrony in the replication sample (*N* = 89) during trait-relevant vs. trait-irrelevant scenes was computed similarly. ISC = inter-subject correlation.

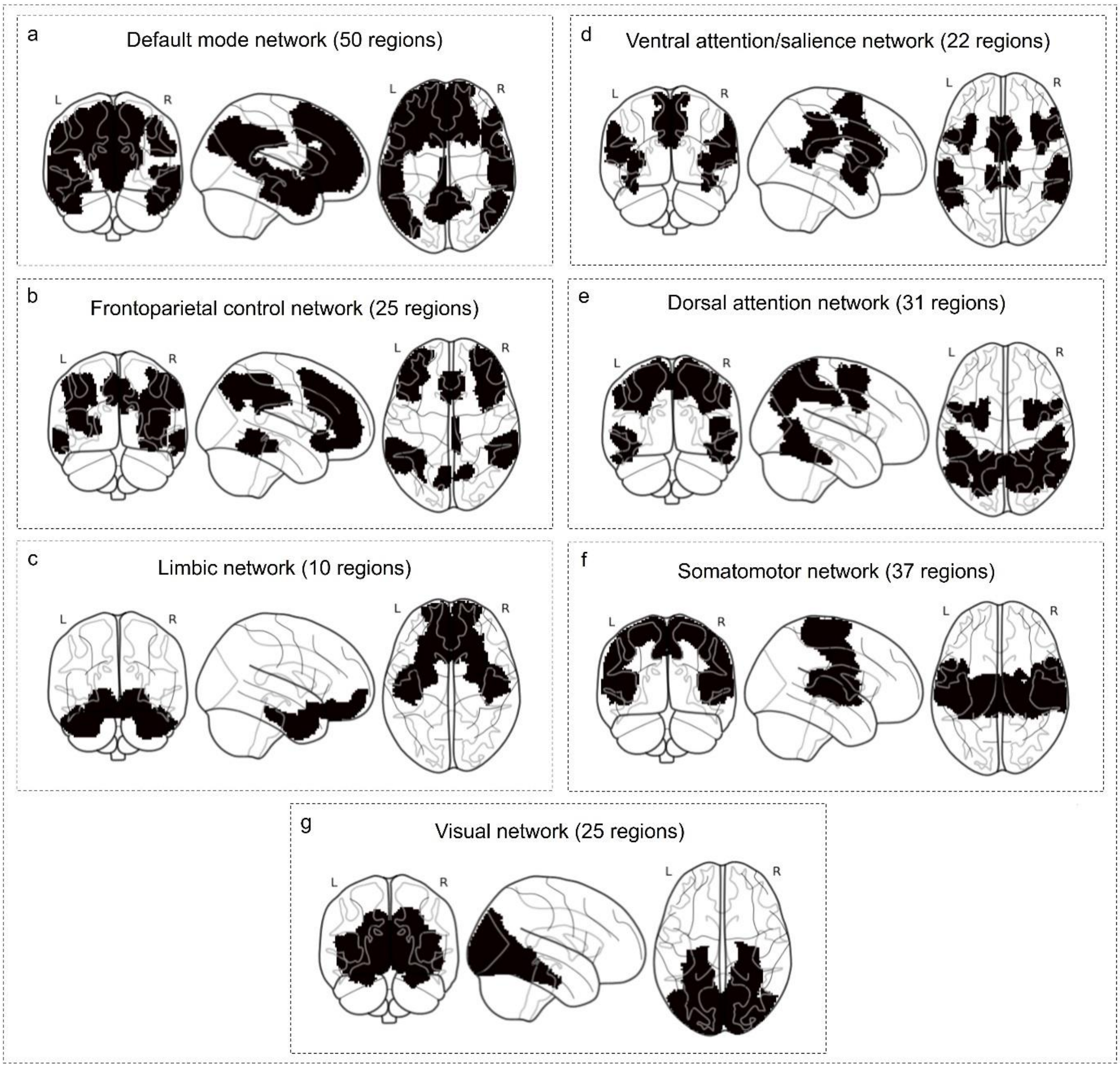


**Supplementary Fig. S6. Visualization of the seven functional Yeo networks.** Each panel depicts the cortical brain regions associated with one of the seven canonical networks defined by Yeo et al.[3]. The number of associated brain regions can be found in parentheses. **(a)** Default mode network. **(b)** Frontoparietal control network. **(c)** Limbic network. **(d)** Ventral attention/salience network. **(e)** Dorsal attention network. **(f)** Somatomotor network. **(g)** Visual network.

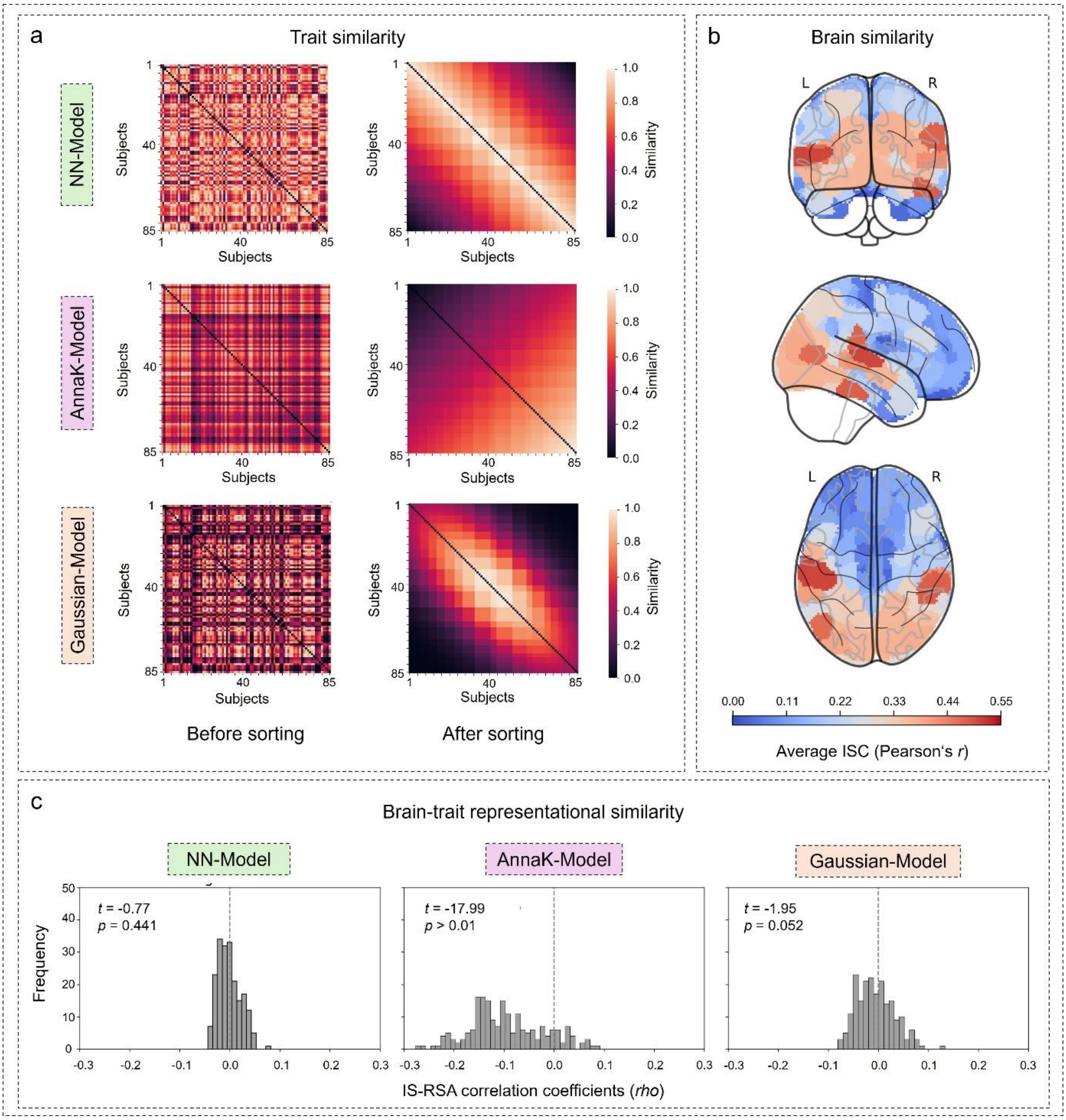


**Supplementary Fig. S7. Inter-subject representational similarity analysis in the replication sample (*N* = 85). (a)** Inter-subject trait similarity in the replication sample is presented for each model implementing different assumptions about the form of brain-trait representational similarity (NN-Model, AnnaK-Model, and Gaussian-Model). In the matrix on the left, subjects are ordered as in the dataset. On the right, the same matrix is shown with subjects sorted by neuroticism rank (from low to high), highlighting the underlying inter-subject similarity structure in neuroticism. **(b)** Brain map reflecting the overall similarity of neural responses across all participant pairs. Specifically, region-specific time courses of brain activity from all scenes were correlated for each pair of participants (ISC). The resulting region-specific ISC correlation coefficients were averaged across all participant pairs to produce a general map illustrating the whole-brain pattern of similarity in brain activity. Higher inter-subject synchrony in brain activity was observed in unimodal sensory regions, whereas multimodal association regions exhibited lower synchrony. **(c)** Frequency distribution of model-specific IS-RSA correlation coefficients computed by comparing the lower triangles of the model-specific *trait* subject-by-subject similarity matrix to each of the 200 brain region-specific *neural* subject-by-subject similarity matrices via Spearman correlation. One-sample *t*-tests were performed to test whether the distribution is significantly shifted from zero, indicating representational brain-trait similarity at the whole-brain level. ISC = inter-subject correlation; IS-RSA = inter-subject representational similarity analysis.

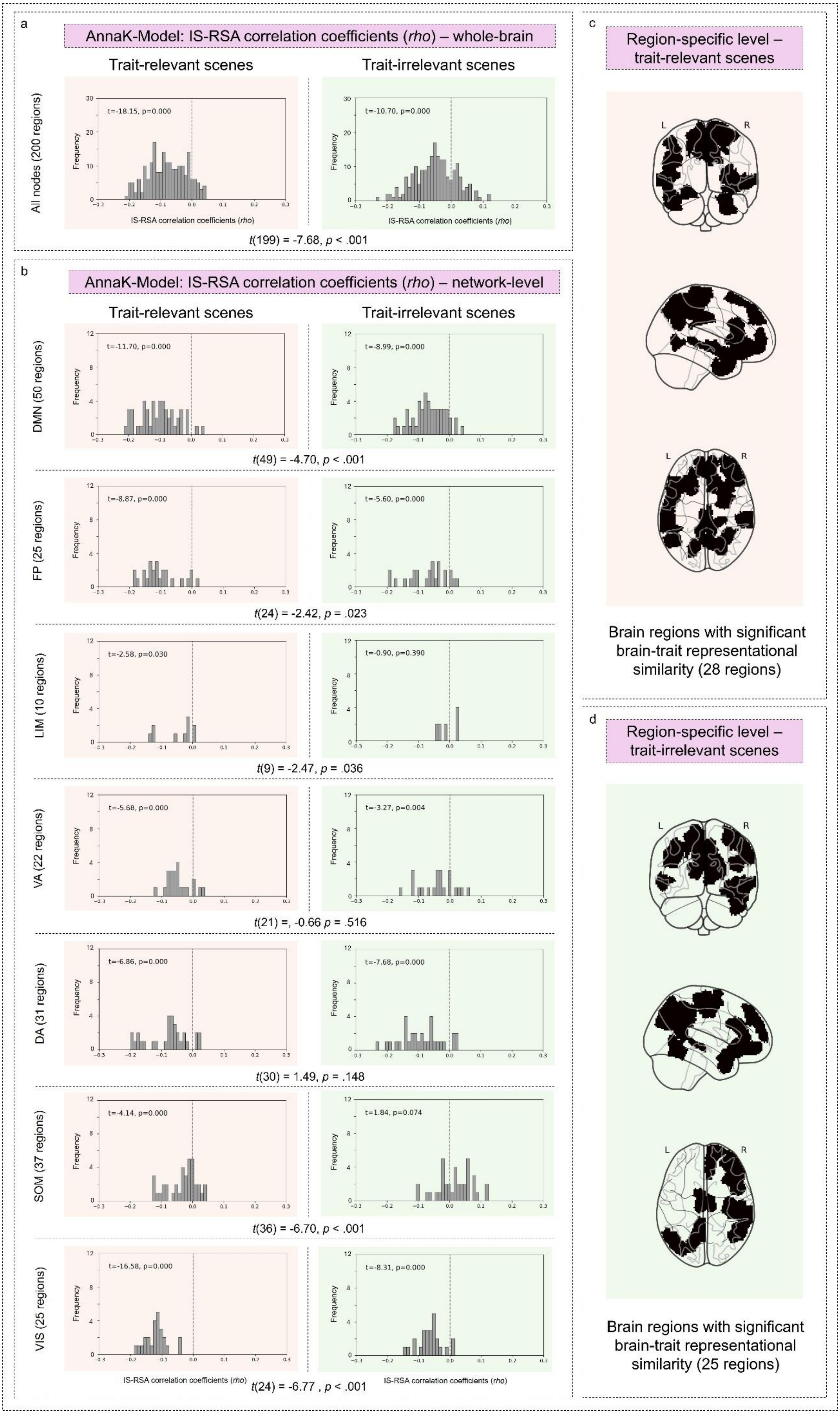
a
AnnaK-Model: IS-RSA correlation coefficients (rho) – whole-brain
Trait-relevant scenes
Trait-irrelevant scenes
All nodes (200 regions)
t=-18.15, p=0.000
t=-10.70, p=0.000
Frequency
IS-RSA correlation coefficients (rho)
t(199) = -7.68, p < .001
b
AnnaK-Model: IS-RSA correlation coefficients (rho) – network-level
Trait-relevant scenes
Trait-irrelevant scenes
DMN (50 regions)
t=-11.70, p=0.000
t=-8.99, p=0.000
t(49) = -4.70, p < .001
FP (25 regions)
t=-8.87, p=0.000
t=-5.60, p=0.000
t(24) = -2.42, p = .023
LIM (10 regions)
t=-2.58, p=0.030
t=-0.90, p=0.390
t(9) = -2.47, p = .036
VA (22 regions)
t=-5.68, p=0.000
t=-3.27, p=0.004
t(21) =, -0.66 p = .516
DA (31 regions)
t=-6.86, p=0.000
t=-7.68, p=0.000
t(30) = 1.49, p = .148
SOM (37 regions)
t=-4.14, p=0.000
t=1.84, p=0.074
t(36) = -6.70, p < .001
VIS (25 regions)
t=-16.58, p=0.000
t=-8.31, p=0.000
IS-RSA correlation coefficients (rho)
t(24) = -6.77 , p < .001
c
Region-specific level – trait-relevant scenes
L
R
Brain regions with significant brain-trait representational similarity (28 regions)
d
Region-specific level – trait-irrelevant scenes
Brain regions with significant brain-trait representational similarity (25 regions)

**Supplementary Fig. S8. Comparison of brain-trait representational similarity under the AnnaK-Model between trait-relevant and trait-irrelevant scenes in the replication sample (*N* = 85) (a)** Frequency distribution of IS-RSA correlation coefficients from the assessment of brain-trait representational similarity during trait-relevant and trait-irrelevant scenes under the AnnaK-Model when considering all 200 brain regions. One-sample t-tests were performed to test whether each distribution was significantly shifted from zero, indicating brain-trait representational similarity at the whole-brain. Further, a paired-sample *t*-test was performed with both sets of 200 IS-RSA correlation coefficients to assess whether brain-trait representational similarity significantly differed between trait-relevant and trait-irrelevant scenes. **(b)** Frequency distributions of IS-RSA correlation coefficients in the replication sample, split by the Yeo seven-network affiliation[3], such that each analysis and respective panel only includes the coefficients corresponding to regions within the respective network. One-sample *t*-tests were performed to assess whether each distribution is significantly shifted from zero, indicating representational brain-trait similarity at the network-level. Further, paired-sample *t*-tests on both respective sets of network-specific IS-RSA correlation coefficients were conducted to test for significant differences in brain-trait representational similarity between trait-relevant and trait-irrelevant scenes. **(c)** Brain regions with significant brain-trait representational similarity during trait-relevant scenes in the replication sample were identified via non-parametric permutation testing (Mantel-test; uncorrected threshold of $p < 0.05$). Corresponding IS-RSA correlation coefficients and *p*-values are shown in Supplementary Table S6. Please note that no brain region remained significant after FDR correction for multiple comparisons. **(d)** Single brain regions with significant brain-trait representational similarity during trait-irrelevant movie scenes in the replication sample (Mantel-test; uncorrected threshold of $p < 0.05$). Corresponding IS-RSA correlation coefficients and *p*-values are shown in Supplementary Table S7. Again, no brain region remained significant after FDR correction. IS-RSA = inter-subject representational similarity analysis; DMN = default mode network; FP = frontoparietal control network; LIM =

limbic network; VA = ventral attention/salience network; DA = dorsal attention network; SOM = somatomotor network; VIS = visual network.

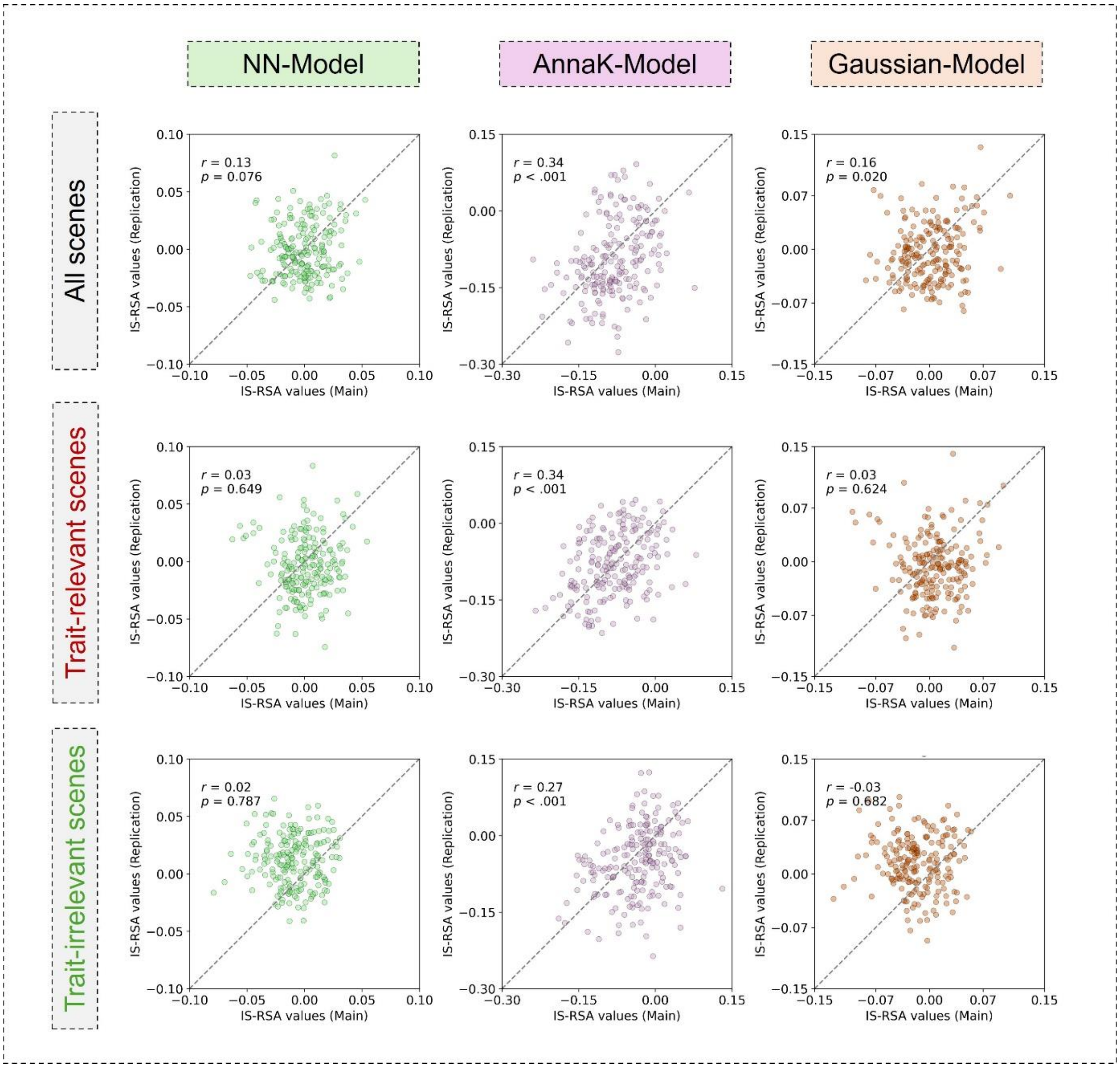


**Supplementary Fig. S9. Comparison of IS-RSA correlation coefficients between the main and replication sample**. Individual scatterplots depict the comparison of IS-RSA correlation coefficients from the main analysis and the replication analysis under each model of trait similarity during all scenes, trait-relevant, and trait-irrelevant scenes. Each dot represents a brain region, and the diagonal depicts the identity line (where x = y). If both analyses gave identical results, all dots would fall on this line. Pearson correlations between model- and region-specific IS-RSA correlation coefficients were computed. Higher values reflect stronger consistency between main and replication results, indicating successful replication. IS-RSA = inter-subject representational similarity analysis.
